# Extruding transcription elongation loops observed in high-resolution single-cell 3D genomes

**DOI:** 10.1101/2023.02.18.529096

**Authors:** Honggui Wu, Jiankun Zhang, Longzhi Tan, X. Sunney Xie

## Abstract

Inside human nuclei, genes are transcribed within a highly packed genome, whose organization is facilitated by cohesin-mediated loop extrusion. However, whether cohesin-mediated loop extrusion participates in transcription is unknown. Here we report that the cohesin-mediated loop extrusion participates in transcription by forming a topoisomerases-regulated transcription elongation loop (TEL), in which cohesin is stalled at the transcription start site (TSS) and gradually extrudes loops asymmetrically until reaching the transcription termination site (TTS). By improving the spatial resolution of single-cell 3D genome mapping to 5 kb with micrococcal nuclease (MNase) in our new single-cell Micro-C (scMicro-C) method, we directly observed the loop expansion of TELs. Furthermore, TEL’s biological function is to ensure high transcriptional burst frequencies by fast re-initiation of RNA Pol II.

**One-Sentence Summary:** Single-cell high-resolution 3D genome structures reveal that cohesin-mediated loop extrusion participates in transcription.

## Main Text

Human genome is hierarchically folded into a myriad of 3D structures (*1-4*), composed of DNA loops at different genomic scales. Indispensable for genome organization, DNA loop extrusion is mediated by the structural maintenance of chromosomes (SMC) complexes— condensin and cohesin, which bind to chromatin and reel flanking DNA into growing loops until the complexes run into roadblocks such as convergently oriented CTCFs (*5-11*). Although the cohesin complex is important for the establishment and maintenance of genome architecture, its depletion only causes modest gene expression changes (*12, 13*). Therefore, we set out to investigate whether cohesin-mediated loop extrusion facilitates transcription involving RNA polymerase II (RNAPII). However, existing technology does not provide enough resolution to address this question.

Chromosome conformation capture (3C or Hi-C) assays have advanced our understanding of 3D genome structures by determining genome-wide contact maps, using restriction enzyme to cut specific sequences to allow the nearby DNA fragments to ligate before conducting whole-genome sequencing (*14, 15*). A “contact” means two DNA sequences that are otherwise separated in the linear (1D) genome but are brought together in the 3D genome. Bulk Hi-C has relatively low resolution due to restriction enzyme having limited cutting sites in the genome. Recently, Micro-C have been developed using micrococcal nuclease (MNase), which enzymatically cuts the genome between two nucleosomes, giving nucleosome-sized fragments for proximity ligation (*16-18*), achieving much higher resolution. However, bulk Hi-C and Micro-C measurements can only measure population-averaged contacts and are unable to distinguish the chromosome positioning and folding among individual cells. To solve this problem, we previously developed Dip-C to determine the 3D genome structure of a single human cell at 20 kb resolution by distinguishing the paternal and maternal alleles based on their single-nucleotide polymorphisms (SNPs) (*19*). In this work, we further improved the spatial resolution of single-cell 3D genome determination by developing single-cell Micro-C (scMicro-C).

### Development of scMicro-C

The previously reported Micro-C chemistry, though offering high spatial resolution, cannot achieve single-cell precision because of substantial loss (over-digestion) of DNA and low ligation efficiency (*16, 17*). To achieve scMicro-C, we made three improvements (Fig. 1A, see Methods). First, we titrated MNase digestion to reduce DNA loss and to produce proper DNA fragments (fig. S2, A and B). Second, we solubilized chromatin with an ionic detergent, sodium dodecyl sulfate (SDS), which dramatically improved ligation efficiency (Fig. 1B). Third, we adopted transposon-based whole-genome amplification method, META (*19*), using Tn5 (*20*), to improve detection of chromatin “contacts.” To confirm such improvements do not compromise data quality, we performed bulk Micro-C (see methods) to generate a high-resolution 3D genome map of human lymphoblastoid cells (GM12878) with 4.4 billion valid contacts from two biological replicates (fig. S1 and table S1). Compared to previously Hi-C data with highest-depth (4.9 billion contacts) (*21*), our Micro-C data detected more chromatin loops (HICCUPS: 20882 vs. 9738) and “stripe” structures (*22*) (Stripenn: 3414 vs. 2722) (fig. S1, E and F).

**Fig. 1.**
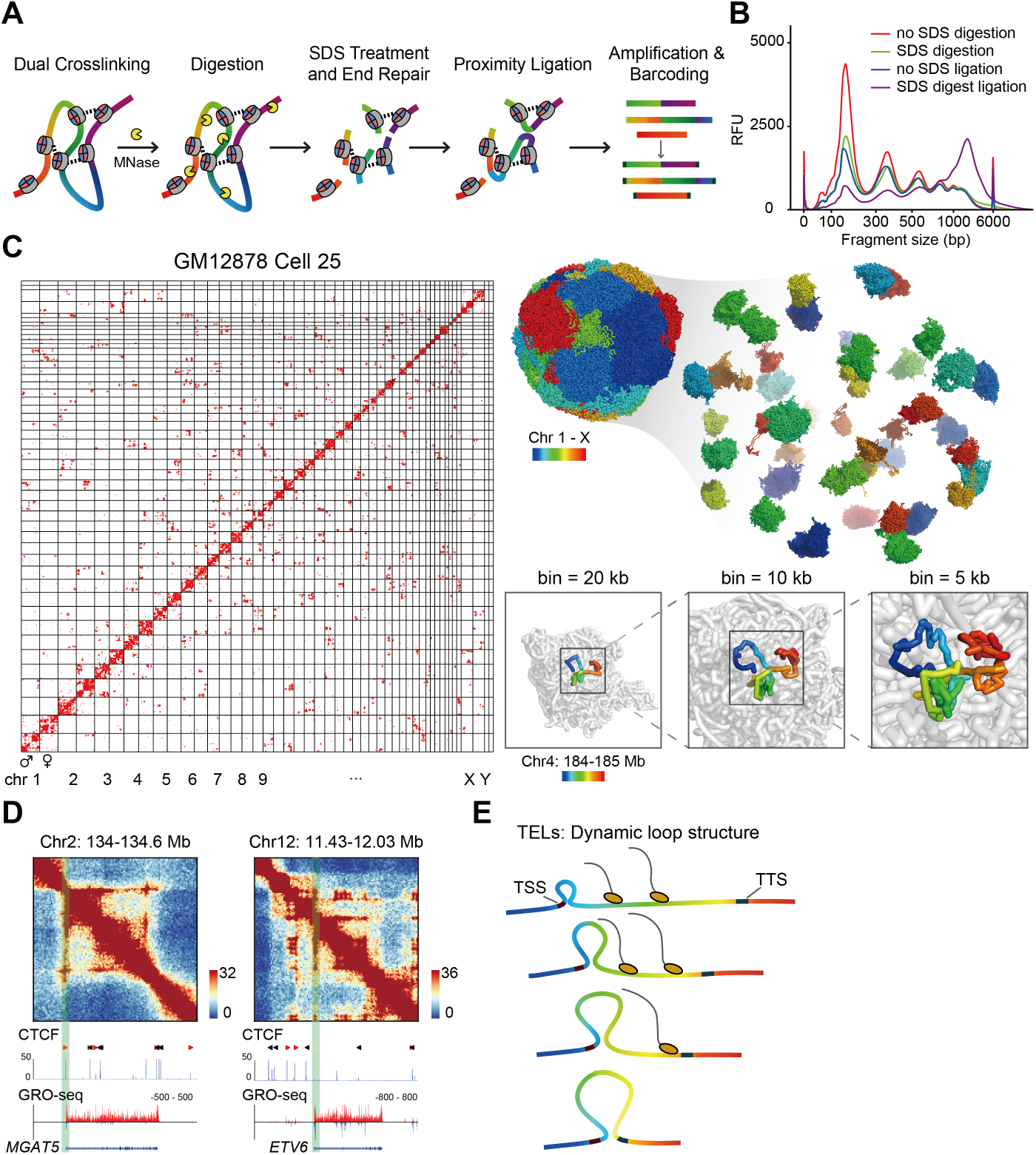
Development of scMicro-C and observation of transcription elongation loops. (**A**) Schematic of scMicro-C procedure. (**B**) Comparison of the ligation efficiency between SDS treatment and the original Micro-C protocol. (**C**) A haplotype-imputed contact map of a representative GM12878 cell (left), and the corresponding 3D genome structure at 5 kb resolution (top right), a selected region (chr4: 184-185 Mb) are shown at 20 kb, 10 kb and 5 kb resolution (bottom right). (**D**) Contact maps of two transcription elongation loops at 5 kb resolution, left: with CTCF binding at TSS, right: no CTCF binding at TSS. CTCF and GRO-seq tracks are shown on the bottom. (**E**) Schematic representing TELs: dynamic loop structure with TSS frequently interacting with downstream loci down to the TTS.

We performed scMicro-C on 340 GM12878 cells, obtaining an average of 1.0 million contacts per cell (SD = 0.5 million, minimum = 0.2 million, maximum = 3.1 million) (fig. S2E and table S2). Compared to our previously developed Dip-C, scMicro-C is more cost-effective and has a higher signal-to-noise ratio (fig. S2, F and G), which is indicated by a much higher contact-to-read ratio (average (± s.d.) of (16.5 ± 5.7)% versus (8.1 ± 2.9)%) and a lower fraction of inter-chromosomal contacts (*16, 17*) (average (± s.d.) = (15.2 ± 6.0)% versus (22.0 ± 1.8)%). Importantly, we proved that scMicro-C preserved the high-resolution and nucleosome occupancy characteristics of Micro-C (fig. S3).

Using our previously developed Dip-C algorithm (*19*), we confirmed that scMicro-C enables the reproducible reconstruction of 3D genome structures in single-cell at 5 kb resolution (Fig. 1C and fig. S4 and S5, see Methods). The pairwise distance matrices show strong concordance with bulk contact maps across different resolutions (Pearson’s r = -0.92 (20 kb), -0.91 (10 kb), -0.89 (5 kb)), demonstrating that the single-cell kilobase-resolution 3D structures are reliable and informative.

### Observation of transcription elongation loops (TELs)

When examining high-resolution bulk contact maps at the scale of single genes, we noted that many active long genes have a line structure on chromatin contact maps, starting from the TSS and ending at the TTS (Fig. 1D and fig. S6), indictive of a dynamic looping structure in individual cells (Fig. 1E). We termed this structure transcription elongation loops (TELs). We stress that these “lines” associated with TELs are different from “architectural stripes,” which are mostly found at TADs boundaries and are formed between two strong CTCF binding sites (*22*). Architectural stripes are usually longer and stronger than TELs (fig. S1F), while TELs can form at genes without CTCF (Fig. 1C and fig. S6A). TELs are as long as genes, whereas architectural stripes are longer than most genes and sometimes appear at genomic regions without genes. We also noted that Zhang et al. reported gene-body-associated domains (*23*), which were observed in highly expressed genes, though the boundary line was not observed perhaps due to the lack of spatial resolution, nor was the mechanism discussed.

We next examined the dependence of TELs on gene length and transcriptional activity. We sorted transcribed genes according to both gene length and RNAPII abundance, and then performed pile-up analysis centered at TSS region (see methods). We found the TELs are most apparent in the longest group (⩾ 50 kb) and strengthened with transcriptional activity (fig. S7A). Such phenomenon is further validated by re-analyzing three published Micro-C datasets (*16, 17*) (fig. S7A and table S3). These results indicated that TELs are closely associated with transcriptional activity and gene length.

We further examined whether active transcription is necessary for the maintenance of TELs by re-analyzing published Micro-C datasets of transcription inhibition by triptolide (TRP) and flavopiridol (FLV) treatments (*16*), which inhibit RNAPII transcription initiation and elongation, respectively. Our re-analysis showed that transcription inhibition by either TRP or FLV treatments both significantly weakens TELs signal (fig. S7B), indicating that transcription is necessary for TELs maintenance of highly expressed genes. It is noteworthy that RNAPII occupancy at TSS is not altered after FLV treatment (*16*), thus, it’s not RNAPII occupancy but transcriptional activity that is needed for TELs maintenance. In contrast, architectural stripes are largely unaffected upon transcription inhibition (fig. S7C). These results further distinguish TELs from architectural stripes.

### Observing TELs with scMicro-C

To confirm the assignment of the line between TSS and TTS in the contact map to the TEL, we used our scMicro-C to visualize the individual 3D structures of a particular gene. We chose Musashi RNA binding protein 2 (*MSI2*) gene (chr17, 428.8 kb, 33.78 FPKM). The *MSI2* gene encodes an RNA-binding protein that are ubiquitously expressed across all tissues, and acts as translational regulator to play a role in the maintenance of stem cells, and is dysregulated in a variety of human cancers (*24*).

For *MSI2*, bulk Micro-C contact map has a TEL line (Fig. 2A, left), consistent with the sum of individual cells’ contact maps (Fig. 2A, right), each giving a corresponding gene structure in a cell (Fig. 2C, upper panel and fig. S8). We selected 6 equally spaced loci between TSS and TTS along the genome (p1–p6) (Fig. 2B) and counted the structures whose 3D distance between the TSS (p0) and another locus is shorter than 3.5 particle radii (∼240 nm). We found that the six loci are evenly populated (Fig. 2C and fig. S9I). Other TEL genes exhibit the same phenomenon (fig. S9, A to F), demonstrating our single-cell 3D structures captured the gradual loop expansion process of TELs (Fig. 2C, upper panel and fig. S8). In contrast, when analyzing a similarly sized genomic region without TELs or stripe structures (chr2: 114.25-115.15 Mb), as expected, we found that the population decreases with larger separation of genomic distance (fig. S9, G to J). These results confirm the existence of enlarging TELs at single-cell and single-gene level.

**Fig. 2.**
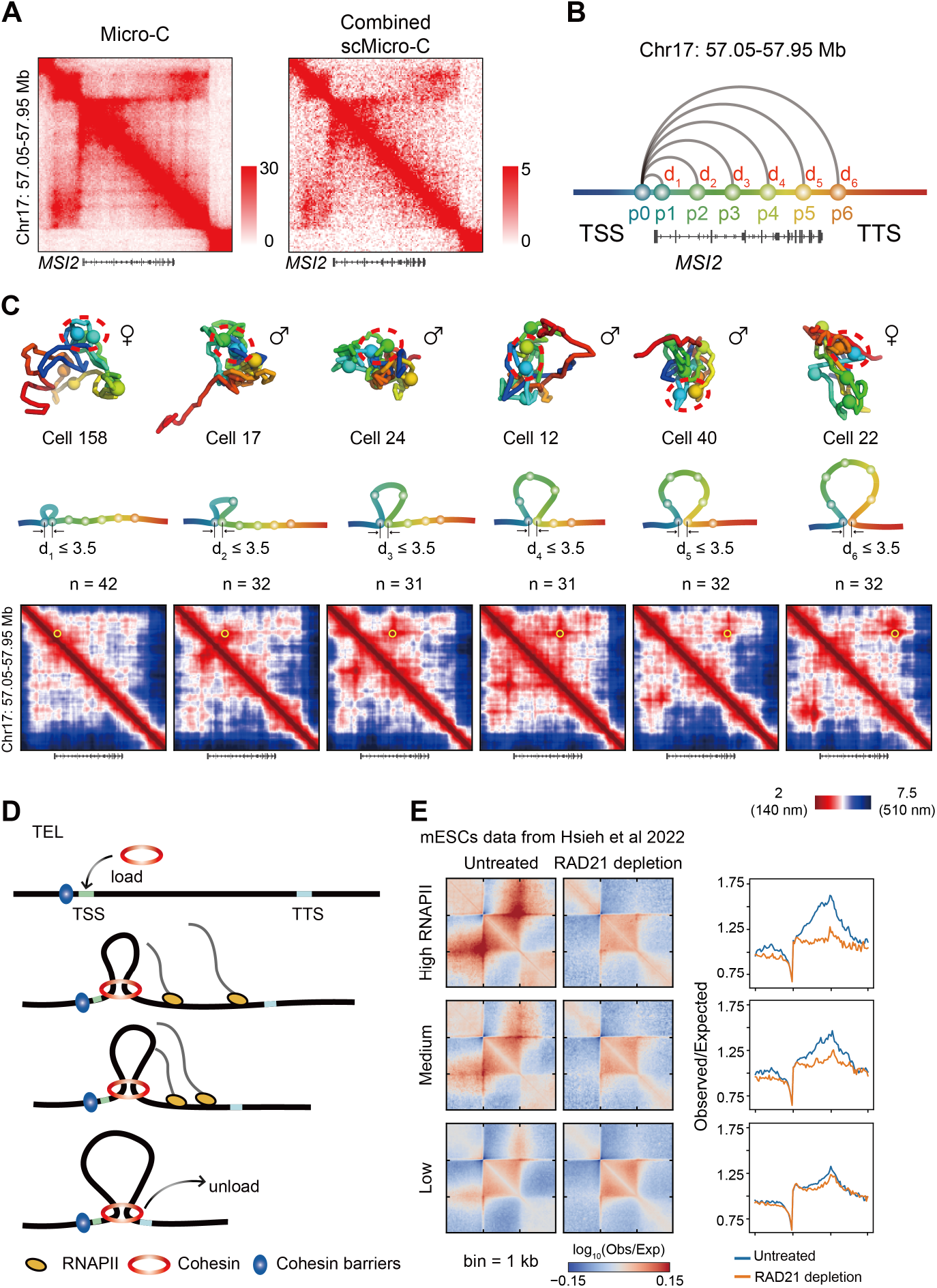
Direct observation of TELs with scMicro-C and cohesin-mediated asymmetric extrusion accounts for TELs. (**A**) Contact maps of bulk Micro-C (left) and combined scMicro-C (right) around MSI2 gene. (**B**) Schematic of loci selection for single-cell 3D structure analysis (p0-p6). (**C**) Top: representative single-cell structures of the sis groups, contacting loci are highlighted. Middle: schematic depiction of the six groups, representing the gradual loop expansion process. Bottom: Average distance matrix of six groups, the number of structures is labeled. (**D**) Proposed model to explain the formation mechanism of TELs, involving cohesin-mediated asymmetric extrusion. (**E**) Cohesin depletion eliminates TELs. Rescaled pile-up analysis of published Micro-C dataset after RAD21 depletion of mESCs at 1 kb resolution (*13*), genes are sorted by transcriptional activity into high (n = 562), medium (n = 475) and low (n = 1986). Only genes ≥ 50 kb are shown. Quantification of TSS interaction enrichment on right.

### TELs are caused by cohesin-mediated asymmetric DNA loop extrusion

Although cohesin depletion doesn’t cause significant changes in gene expression (*12*), literature showed that cohesin may be involved in transcription. First, according to ChIP-seq, though most cohesin colocalizes with CTCF, which serves as a cohesin barrier at TADs boundary, roughly 1/4 of cohesin binds to active TSS, indicating active promoters could act as cohesin barriers in mouse and human (*25, 26*). Second, NIPBL, a cohesin loading factor, is enriched at promoters of active genes (*27*), indicating that cohesin prefers to load at TSSs. Third, in the absence of CTCF and cohesin unloading factor WAPL, cohesin accumulates at the 3’ end of some active genes (*25*), implying that cohesin tends to unload at TTSs. As shown in the Fig. 2D, we propose a model to explain the formation of TELs involving cohesin-mediated asymmetric loop extrusion: cohesin is preferentially loaded near active gene promoters, but is stalled at the TSS by a barrier, and thus can only extrude the gene body side until reaching and detaching at the TTS, thus forming the TEL.

We verified that cohesin is indeed involved in TEL formation by re-analyzing a published high-resolution bulk Micro-C dataset after acute cohesin depletion (*13*). In untreated cells, we observed strong TEL signals in highly transcribed genes (top panel) compared to other genes (middle and bottom panel). In contrast, these signals are completely eliminated after acute cohesin depletion (enrichment scores drop from 1.50 to 1.13) (Fig. 2C and fig. S10A). Therefore, TELs are dependent on cohesin-mediated loop extrusion.

### Transcription of TEL-associated genes is more sensitive to topoisomerase inhibition

During transcription, RNAPII generates positive supercoiling in the forward direction and negative supercoiling in the backward direction (*28*), which are both released by DNA topoisomerases (*29*). In humans, there are mainly two types of topoisomerases, type IB (TOP1) and type IIA (TOP2A and TOP2B), which remove both positive and negative supercoiling. TOP1 cuts only one strand of double-stranded DNA and re-ligate it after supercoil release, and is associated with transcribing RNAPII, exhibiting catalytic activity at gene body and 3’ end of active genes (*30*). On the other hand, TOP2 introduces double-strand DNA breaks at where two DNA duplexes cross, and rejoins the two ends after supercoil release. TOP2 activity is concentrated at promoter-proximal regions, depending on the expression level and gene length (*31*). During their catalytic cycles, transient topoisomerase-DNA covalent complexes (TOPccs) can be inhibited by many small molecules (*32, 33*), which we used below to study this process.

We first evaluated the association between topoisomerase and TELs. A previous study found that TOP1 inhibition downregulates the expression of long genes in neurons (*34*). To identify genes affected by topoisomerase inhibition, we performed RNA-seq on GM12878 cells treated with TOP1 inhibitor topotecan (TPT) or TOP2 inhibitor etoposide (ETO) by following the previously documented experimental condition (*34*) (fig. S11, B to D) (see Methods). To examine the effect of topoisomerase inhibition on cell cycle, we performed cell cycle analysis and found only a slight increase of cells in the S phase and a slight decrease of cells in the M phase (fig. S11A), confirming that the inhibition experiment is valid.

As shown in fig. S12 E, among differentially expressed genes, 989 genes are downregulated (68% in ETO and 29% in TPT) and 1791 genes are upregulated (60% in ETO and 46% in TPT) by either TPT or ETO treatment (false-discovery rate (FDR) < 0.01, fold change > 2). Downregulated genes are biased towards long genes (Fig. 3A), especially for TOP1 inhibition, exhibiting strong negative correlation between gene length and gene expression fold change (Pearson’s r = -0.476) (fig. S11F). We found that downregulated genes are significantly enriched in two categories: housekeeping genes and immune-related (B cell-specific) genes (Fig. 3B and fig. S11, G and H). These results reveal that TOP1 and TOP2 are important for maintaining expression of long and highly expressed genes.

**Fig. 3.**
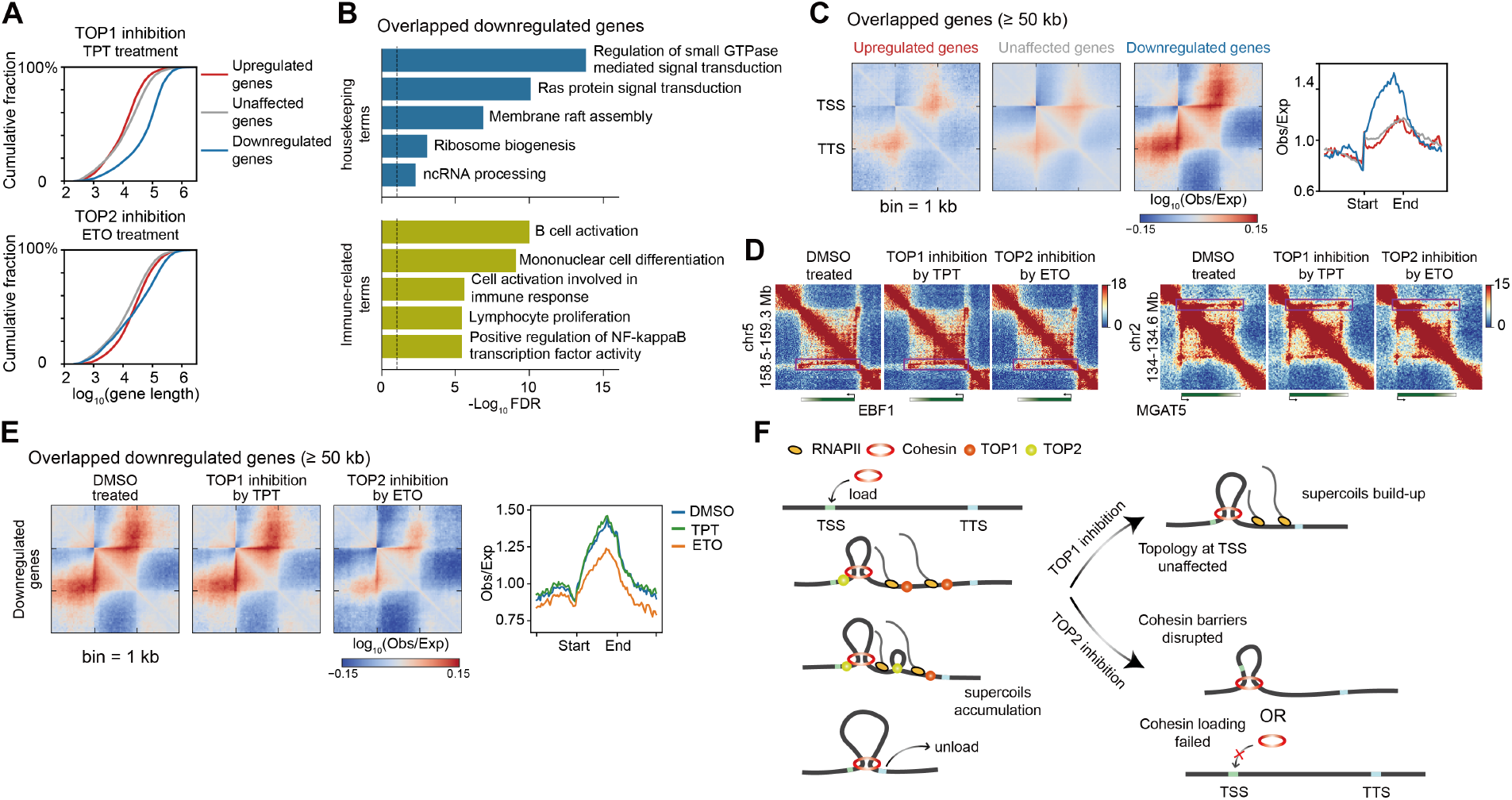
Different involvement of topoisomerase I and II involved in TELs. (**A**) Cumulative curves of gene length of genes affected by TOP1 inhibition (top) and TOP2 inhibition (bottom). (**B**) Gene Ontology (GO) terms of genes downregulated by either TOP1 or TOP2 inhibition, these terms are grouped into two categories: housekeeping and B cell-specific term. (**C)** Rescale pile-up analysis of genes ≥ 50 kb at 1 kb resolution. Overlapped genes between TOP1 and TOP2 inhibition are used, upregulated (n = 416), unaffected (n = 3254) and downregulated (n = 583). Quantification is shown. (**D**) TELs are weakened after TOP2 inhibition. Contact map of two TELs for DMSO control, TOP1 (TPT) and TOP2 (ETO) inhibition Micro-C datasets at 5 kb resolution. The quantification of TSS interaction enrichment is shown on right. (**E**) Rescaled pile-up analysis of genes ≥ 50 kb at 1 kb of DMSO control, TOP1 and TOP2 inhibition Micro-C datasets, overlapped downregulated genes upon TOP1 and TOP2 inhibition are used. Quantification is shown on right. (F) Proposed model to explain how TOP1 and TOP2 are differently involved in TELs maintenance.

We next examined whether topoisomerase-sensitive genes have TELs and found that genes downregulated by topoisomerase inhibition, corresponding to about 1/5 of all expressed long genes, exhibited strong TEL signals (Fig. 3C and fig. S12). We noted that genes specifically downregulated by TOP1 inhibition showed weaker TEL signals than genes downregulated by TOP2 (fig. S12), indicating that TOP1 is required for the proper expression of many long genes without TEL due to the length-dependent effect of supercoiling build-up (*34*). We speculated that TEL-associated genes are more likely to accumulate torsional stress due to that cohesin-mediated asymmetric extrusion restricts the dissipation of transcription-induced supercoiling (Fig. 3F, left panel), which necessitates topoisomerase. Thus, genes with TELs, corresponding to about 1/5 of all expressed long genes, are observed to be facilitated by topoisomerases.

### Distinct roles of Topoisomerases 1 and 2 in TELs

To investigate the effect of TOP1 and TOP2 on TELs, we generated high-resolution Micro-C contact maps for GM12878 cells treated with TOP1 inhibitor (TPT), TOP2 inhibitor (ETO) and control (DMSO), and obtained 2.5, 2.6 and 2.4 billion valid contacts from two independent biological replicates (table S1). We found that TEL signals are greatly weakened upon TOP2 inhibition, while TEL signals are slightly strengthened upon TOP1 inhibition (Fig. 3, D and E and fig. S13). In contrast, we found that architectural stripes are largely unaffected by either TOP1 or TOP2 inhibition (fig. S14). These results indicated that topoisomerases are involved in TELs.

We now explained the mechanism underlying the different involvement of TOP1 and TOP2 in the establishment or maintenance of TELs. Upon TOP1 inhibition, elongating RNAPII are stalled due to supercoiling build-up at gene body (*34*), which further interferes cohesin extrusion. However, TOP1 does not exhibit activity at TSS region (fig. S13E), preserving the barrier of cohesin at TSS and thus explaining why TELs are not disrupted by TOP1 inhibition (Fig. 3F, top right). It was previously known that TOP2 maintains the negative supercoiling at TSS (*35*). Upon TOP2 inhibition, negative supercoiling at TSS is disrupted, causing two possible scenarios: loss of barrier at TSS, and/or failure of cohesin loading at TSS. Both scenarios would lead to abolishment of TELs (Fig. 3F, bottom right).

### TELs assure high transcriptional burst frequency

We now answer the question of what the physiological significance of TELs is. Transcription of gene is bursting even for highly expressed genes in both prokaryotes and eukaryotes because of single-molecule nature of DNA (*36-38*), which have been extensively studied by real-time imaging on single-cell basis and single-cell transcriptomics genome-wide. In bacteria, transcriptional bursting was shown to be due to positive supercoiling buildup (*38*). In both eukaryotic and prokaryotic cell, transcription can be described by the two state model (*39*), where transcription stochastically switches between “on” and “off” states (Fig. 4a). To measure transcriptome-wide transcriptional burst kinetics, we fitted our full-length single-cell RNA-seq data (*40*) to determine the burst frequencies (k_on_) and burst sizes (u/k_off_) (fig. S15, A to C). We noted that inferred k_on_ values were much smaller than k_off_ values (fig. S15D), confirming that off states dominate the transcription burst cycle (*41*), which we assume is primarily caused by transcription re-initiation at least for long genes.

**Fig. 4.**
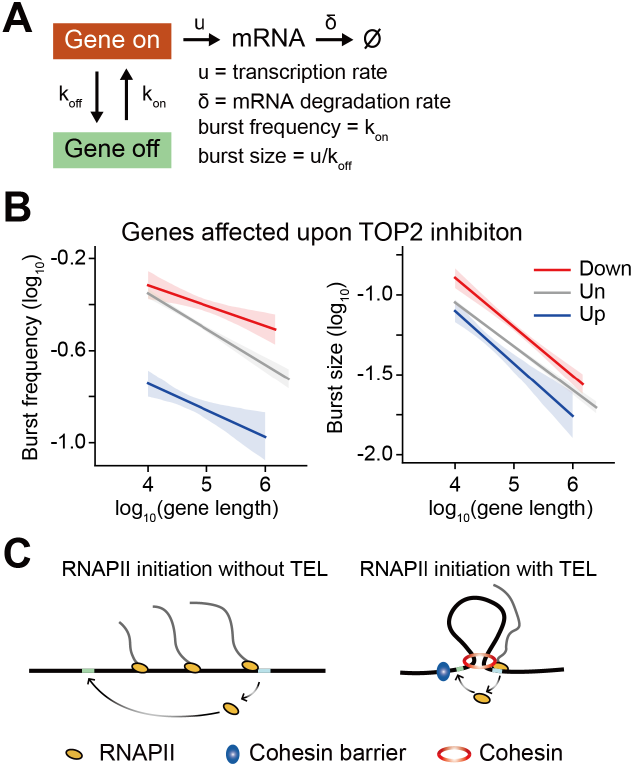
Transcription elongation loops ensure high transcriptional burst frequency. (**A**) Two-state transcription model, genes stochastically switch between on and off at rate of k_off_ and k_on_. (**B**) Burst frequency and burst size as a function of gene length, genes are grouped based on TOP2 inhibition RNA-seq. (**C**) Model to clarify how TELs enable faster RNAPII recycling.

We found that genes with TELs that are downregulated by TOP2 inhibition have higher burst frequencies than other genes (Fig. 4B, left and fig. S15, G to I). In contrast, the difference of the burst size between TEL-associated genes and other genes are not obvious (Fig. 4B, right and fig. S15, G to I). These results demonstrated that cohesin-mediated TELs guarantee high transcriptional burst frequency of highly expressed long genes. In fact, this could explain why short-term cohesin depletion by degron has a little effect on nascent transcription level (*12*), because it only spans about 1-2 burst cycle (2-3 hrs).

We therefore propose a model to explain how TELs enable high transcriptional burst frequency (Fig. 4D). For long genes without TELs, dissociated RNAPII takes a long time to search for and rebind to promoters for the next round of transcription (Fig. 4D, left). In contrast, genes with TELs have their TSS close to their TTS in 3D, which allows dissociated RNAPII to take less time to rebind to promoters (Fig. 4D, right), thus enabling faster transcription reinitiation. Consistent with this model, we observed that TOP2 inhibition-downregulated genes have stronger enrichment for proteins belonging to transcription initiation complexes (PLOR2A, MED1, TBP and TAF1) at their TTS regions, demonstrating TTSs form strong interactions with their TSSs (fig. S16A).

## Discussion

Our understanding of cohesin-mediated loop extrusion and its role in genome organization has been greatly advanced by recent *in vitro* single-molecule (*9, 10*) and *in vivo* inducible protein degradation (*12*) studies. In addition to its role in genome organization, cohesin-mediated loop extrusion has also been proposed to explain the immunoglobulin gene V(D)J recombination (*42*), antibody class switching (*43*) and the repair of DNA double-strand breaks (DSB) (*44*). However, the role of cohesin-mediated loop extrusion in transcription, in any, has not been revealed. Our study proved that cohesin participates in the transcription of highly expressed long genes by forming TELs. Specifically, our high-resolution scMicro-C directly captured the progressive loop expansion of TELs, which has not been observed by previous bulk measurements and low-resolution scHi-C.

Many long genes are involved in neuronal functions and specifically expressed in the brain, their dysregulation is associated with neurological disorders (such as autism spectrum disorder and Rett syndrome) and aging (*34, 45*). We note that there is circumstantial evidence in the literature pointing to TEL. For example, a recent report showed that STAG2 (an alternative subunit of cohesin) knockout downregulated genes form promoter anchored stripes (“P-stripe”) in mouse oligodendrocytes (*46*). Our study provided definitive proof of TELs for highly expressed long genes. We note that we cannot rule out the possibility of the existence of TEL in short genes, which cannot be resolved by current Micro-C experiments. However, the biological significance of TEL seems to be more obvious for long genes than for short genes.

Although we experimentally observed the TELs are stalled at TSS (Fig. 2C), the nature of barrier for cohesin at TSS is different from the conventional choice of CTCF. In fact, only a small fraction of genes has promoter-bound CTCF (19.6% within ± 5 kb of TSS in GM12878, fig. S16, B and C), thus CTCF is not the universal TSS barrier. Besides CTCF, other chromatin-bound proteins (e.g., dCas9 (*42*), minichromosome maintenance (MCM) complex (*47*), transcription factors (*48*)) and specific non-canonical DNA structures (e.g., R-loops (*49*), replication folks (*50*)) have been implied as potential cohesin barriers, demonstrating that the barrier doesn’t have to be a DNA-binding protein. In this study, we showed that inactivation of RNAPII (fig. S7B) or TOP2 (Fig. 3E) both disrupt TELs, which indicates that the cohesin barrier at TSS is transcription- and DNA supercoiling-associated, implying it is transcription-induced negative supercoiling.

Transcription, being such a fundamental process in biology, involve complicated coordination with companying dynamical processes, such as positive and negative supercoiling, in the crowded 3D genome space. The discovery of TEL involvement in transcription is thus fundamental importance to the understanding of transcription at molecular and cellular level.

## Acknowledgments

We thank the staff at the Peking University High-throughput Sequencing Center (HTSC) for help in next generation sequencing and flow cytometry; Beijing Berry Genomics for help in next-generation sequencing; We thank Professors Xiong Ji and Xuefei Zhang at Peking University for helpful discussions and suggestions.

## Author contributions

H. W., L.T. and X.S.X. designed the experiments; H.W. performed the experiments; J.Z. analyzed the data; H. W., J.Z., L.T. and X.S.X. wrote the manuscript.

## Data and materials availability

Raw sequencing data generated during this study is deposited at the Sequence Read Archive with accession number PRJNA788160. The code used in this study is publicly available at GitHub (https://github.com/tanlongzhi/dip-c and https://github.com/lh3/hickit).

## Supplementary Materials

Materials and Methods

Supplementary Text

Figs. S1 to S16

Tables S1 to S3

References (*51-52*)

