## Supplementary material for "Extruding transcription elongation loops observed in high-resolution single-cell 3D genomes": scMicro-C_Supplementary_Materials.pdf

**The PDF file includes:**

Materials and Methods
Supplementary Text
Figs. S1 to S16
References (51-52)

### Materials and Methods

#### *Cell culture*

GM12878 lymphoblastoid cells were grown in RPMI1640 medium (Gibco, Thermo Fisher Scientific, Cat. # 11875093) supplemented with 15% FBS (Gibco, #10091148) and 1% Pen/Strep (Gibco, #15140122), maintained at 37°C with 5% CO2 at recommended density. Upon harvest, cells were spun down and washed once with ice-cold PBS.

*Topoisomerase inhibition.* For topoisomerase inhibition experiments, we use both TOP1 and TOP2 inhibitors. First dissolved inhibitors in DMSO to make stocks (3 mM for topotecan (APExBIO, B4982), 50 mM for etoposide (MCE, HY13629)), then added 10 μL to 10 mL culture medium to a final concentration of 3 μM for topotecan, 50 μM for etoposide. For control, equal volume of DMSO were added to culture medium and cells were cultured for 8 hours.

#### *Bulk RNA-seq*

RNA were extracted with a Trizol-based RNA isolation protocol. Briefly, at least 1x 10^6 cells were harvested and washed twice with ice-cold PBS, then added 1 mL Trizol, incubated at room temperature for 5 min. At the end of incubation, 0.2 mL chloroform were added, vortexed at vigorously for 15 s, then incubated at room temperature for another 5 min. Followed by centrifuging at 10,000 rpm for 5 min at 4°C. Transferred upper aqueous phase carefully without disturbing the interphase into a fresh tube, added 0.5 mL of isopropyl alcohol to supernatant and incubated at room temperature for 10 min. Centrifuged at 14,000 rpm for 20 min. Discarded supernatant, added 1 mL 75% ethanol, re-centrifuged at 9,500 rmp for 5 min. Air-dry, added nuclease-free water to elute.

*RNA-seq library preparation.* 1 μg total RNA were first captured with mRNA capture beads (Vazyme, #N401), then submitted to RNA-seq library preparation kit (Vazyme, NR605). Each sample prepared 2 replicates, and sequenced at Illumina NovaSeq 6000 platform, each sample sequenced to 100 million reads.

#### *Smart-seq2*

Smart-seq2 was performed according to previous described (*51*) with small modifications. Viable cells were sorted to 96-well plates containing 2 μL lysis buffer (0.15% Triton X-100, 1 mM dNTP, 1 μM oligo-dT, 1 U/μL RNase inhibitor), incubate at 72°C for 10 min. Then 3 μL reverse transcription mix (1x first strand buffer, 1 mM GTP, 5 mM DTT, 1 M Betaine, 6 mM MgCl2, 1 μL TSO, 10 U/μL SSII, 1 U/μL RNase inhibitor) was added to each well, incubated as 42°C for 90 min, 10 cycles [50°C, 2 min; 42°C, 2 min], 70°C, 5 min. After RT, 15 μL amplification mix was added to each well, incubated as 98°C, 3 min, 21 cycles [98°C, 20 s; 65°C, 30 s; 72°C, 4 min], 72°C, 5 min, hold at 4°C. Purified with 0.7x AMPure XP beads, then use home-made Nextera transposome for library preparation.

#### *Modified bulk Micro-C*

Bulk Micro-C protocol was adopted from published Micro-C protocol for mammalian cells (*16, 17*) with several modifications. Briefly, harvested cells were crosslinked with freshly made 1% PFA followed by 3 mM DSG, then cells were lysed and titrated to test the appropriate MNase concentration (fig. S2B). Before end repair, we added 50 μL 0.3% SDS and incubated at 62°C for 10 min, then quenched by 50 μL 3% Triton X-100 and incubated at 37°C for 15 min. Then nuclei were end repaired and biotin labeled, followed by *in situ* ligation with T4 ligase. Before DNA extraction, nuclei were incubated with exonuclease III (NEB, #M0206) to remove un-ligated ends. Extracted DNA were either sonicated to 300 bp or select di-nucleosome fragments for library preparation. Then performed biotin pull down and adaptor ligation for sequencing. We generated two replicates, one with sonication procedure, one with di-nucleosome selection procedure.

***Single-cell Micro-C protocol***

Our protocol was modified from mammalian cell Micro-C protocols with three key modifications. First, we optimize Micrococcal nuclease (MNase) digestion level to produce longer DNA pieces and reduce DNA loss, to ensure 40%-50% mononucleosomes and 20-25% di-nucleosomes (average 500 bp) instead of 80-90% mononucleosomes and 10-20% di-nucleosome (average 200 bp) as original Micro-C suggested, because over-digestion produces too short fragments, which hampers transposon-based whole genome amplification procedures. Second, we add an ionic detergent (SDS) to solubilize chromatin between MNase digestion and end repair step, which both preserves Micro-C characteristic nucleosome-resolution chromatin interactions (as suggested by our bulk data) and dramatically increases ligation efficiency (fig. S2C), further increase the length of the final product from ~450 bp (without SDS) to ~1300 bp, which is critical for single-cell WGA. Third, we use our state-of-the-art WGA method, META, to further increase contact detection efficiency in single cell. We omit all biotin-related steps to maximize the number of contacts detected per cell as described in Dip-C procedure (*19*).

*Cell Crosslinking.* Cells were fixed with 1% PFA (EMS, 15714) at room temperature with rotation. PFA was quenched by the addition of 2 M Tris-HCl pH 7.5 to a final concentration of 0.75 M and incubated at room temperature for 5 min. Then wash twice with ice-cold 1 x PBS supplemented with 1 x BSA (centrifugation: 3000g, 5 min). After crosslinking, some pellets appear and disappear after twice washing. Then cells were further fixed in 3mM DSG in PBS, and incubated at room temperature for 45 min. DSG was quenched by the addition of 2 M Tris-HCl pH 7.5 to a final concentration of 0.75 M and incubated at room temperature for 5 min. After fixation, cells were washed twice with ice-cold 1 x PBS supplemented with 1 x BSA (centrifugation: 3000g, 5 min). The pellets were stored at -80°C.

*MNase Digestion.* 1 million cells were permeabilized with 100 μL Micro-C buffer1 (50 mM NaCl, 10 mM Tris-HCl pH 7.5, 5 mM $MgCl_2$, 1 mM $CaCl_2$, 0.2% IGEPAL CA630, 1 x PIC), and incubated on ice for 20 min. Then resuspend the cell pellet in 100 μL Micro-C buffer1, then titrate to find the appropriate amounts of MNase (NEB M0247S) to digest chromatin to 40-50% mononucleosome and 20-25% di-nucleosomes (fig. S2B), and incubate at 37 ° C for 10 min. Add 0.5 M EGTA to the final concentration of 4 mM to stop the reaction, omitting the heat inactivation step.

*End repair.* Resuspend cell pellets in 50 μL 0.5% SDS, incubate at 62°C for 10 min, then add 170 μL 1.5% Triton X-100, incubate at 37°C for 15 min. Then wash once with 100 μL Micro-C buffer2 (50 mM NaCl, 10 mM Tris-HCl pH 7.5, 10 mM MgCl2). End chewing was executed with two steps: first, resuspend the cell pellet in 45 μL End repair buffer1 (1 x NEBuffer 2.1, 2 mM ATP, 5 mM DTT, 2.5 μL 10 U/μL T4 PNK), then incubate at 37°C for 15 min with gentle vortex to add 5' phosphate and remove 3' phosphoryl groups; next, add 5 μL 5 U/μL Klenow fragment, then incubate at 37°C for 15 min with gentle vortex to remove 3' overhangs. Blunt end repair was performed by the addition of 25 μL End repair buffer2 (200 mM dNTP/each, 1 x T4 Ligase buffer, 100 μg/mL BSA), incubated at room temperature for 45 min. Wash once with 1 mL Micro-C buffer 3 (50 mM Tris-HCl pH7.5, 10 mM MgCl2).

*Proximity Ligation.* Ligation was performed by the addition of 250 μL ligation mix (1 x T4 Ligase buffer, 100 μg/mL BSA, 20U/μL T4 Ligase), and rotating at room temperature for 2.5 hours. Then the cell pellet was stained with DAPI/7-AAD, DAPI/7-AAD positive nuclei were sorted with flow cytometry.

***Plate-based single-cell amplification***

Nuclei were sorted to 96-well PCR plates containing 2 μL lysis buffer (10 mM Tris pH 8.0, 20 mM NaCl, 1 mM EDTA, 0.1% Triton X-100, 500 nM Carrier ssDNA, 1.5 mg/mL QIAGEN protease), then lysed nuclei with procedure (50°C, 1 hr, 65°C, 1 hr, 70°C, 15 min). After lysis, nuclei could be stored at -80C for several months. Lysed nuclei were first transposed by the addition of a 6 μL transposition mix ((leading to a final concentration of 10 mM TAPS pH 8.5, 5 mM MgCl2, 8% PEG 8000, 0.3 nM META transposome dimer), and incubated at 55°C for 10 min. META transposome was assembled as previously described (*19*). Transposition was stopped by the addition of a 2 μL stop mix (250 mM NaCl, 37.5 mM EDTA, 2 mg/mL QIAGEN protease) and incubation at 50°C for 30 min, 70°C for 15min. The barcoding strategy is the same as previously described (*52*). Then transposed DNA were amplified by the addition of 15 μL preamplification mix (12.5 μL 2 x Q5 master mix, 0.8 μL 50 μM META16 primer mix, 0.5 μL 100 mM MgCl2, 1.2 μL H2O) and incubated at 72°C for 5 min, 98°C for 30 s, 12 cycles of [98°C for 10 s, 62°C for 30 s, 72°C, 2 min], 65°C, 5 min. Next, add 0.8 μL 50 μM indexed META16-ADP1 primer and 0.8 μL 50 μM META16-ADP2 primer to generate a 12 x 8 cell barcode combinations for each 96-well plates, and incubate at 98°C for 30 s, 3 cycles of [98°C for 10 s, 62°C for 30 s, 72°C, 2 min], 65°C, 5 min. After cell barcoding, a whole plate was pooled together for purification with ZYMO DCC5.

***Library preparation***

120 ng (10 μL) of the purified amplicon was used for each plate for library preparation. Add 40 μL PCR mix (25 μL 2x Q5 Master mix, 5 μL NEBNext index primer i5 (E7600S), and 5 μL NEBNext index primer i7 (E7600S), 0.05 μL 100 mM MgCl2) and incubated at 98°C, 30 s, 2 cycles of [98°C, 10 s, 68°C, 30 s, 72°C, 2 min], 72°C, 5 min. Then purified with 0.8 x SPRI beads to remove < 300 bp fragments.

***Bulk RNA-seq analysis***

Bulk RNA-seq data were mapped to the human reference genome GRCh38 using STAR and the associated gene annotation file was downloaded from GenCode (v34). We used RSEM to count the number of mRNA fragments. Differential expression analyses were then performed using the DESeq2 package. For differentially expressed genes, gene ontology (GO) and gene set enrichment analysis (GSEA) was performed using the clusterProfiler package.

***Single-cell RNA-seq analysis***

Single-cell RNA-seq data were processed similarly to bulk RNA-seq data but using the "--single-cell-prior" option with RSEM package. To remove lowly expressed genes, we only retained genes with a count > 50 in at least 5 cells. The stochastic transcription burst can be described with a two-state model where $k_{on}$ is the rate at which the genes transition from the "off" to the "on" state; $k_{off}$ is the rate at which the genes transition from "on" to "off" state; s is the transcribing rate. The expression count matrix generated above was then used to estimate the transcription burst frequency ($k_{on}$) and size ($s/k_{off}$) using the PoissonBeta algorithm. A bootstrap-based goodness-of-fit test was performed and genes with a bootstrap P-value < 0.1 were filtered out. This approach yielded identifiable kinetic parameters for 10229 genes. Linear regression models were fitted using the seaborn lmplot function.

***Analysis of bulk Micro-C data***

*Generation of contact maps.* Bulk Micro-C datasets were processed using the distiller pipeline (https://github.com/open2c/distiller-nf). Raw FASTQ files were mapped to the human reference genome assembly GRCh38 using the BWA-MEM. Pairs were extracted from the mapped reads using the pairtools package (https://github.com/open2c/pairtools). PCR duplicates were then filtered out and only pairs with MAPQ > 20 were kept. Contact matrices in the .mcool and .hic format were generated and balanced using the cooler (https://github.com/open2c/cooler) and Juicer package (https://github.com/aidenlab/juicer/).

*Contact scaling curves.* We used the cooltools expected-cis and logbin-expected functions (https://github.com/open2c/cooltools) to calculate the normalized contact probability as a function of genomic separation within chromosome arms and the scaling derivatives of the curve on contact matrices at 1-kb resolution. The contact scaling curves using unbinned short-range contacts (contact distance < 10kb) were calculated separately for each contact orientation (IN-OUT, IN-IN, and OUT-OUT) using the cooltools compute_scaling function.

*A/B compartments.* We used the cooltools eigs-cis function to calculate A/B compartments at 500-kb and 100-kb resolution.

*Insulation scores.* The cooltools insulation function was used to calculate the insulation scores at 10-kb resolution (with a window of 100kb). Genomic loci with boundary strength > 0.2 were considered insulation boundaries and used for downstream analyses. We used the bedtools intersect function to compare two lists of insulation boundaries and a 10-kb offset on each side was tolerated when performing this intersection. Average profiles of insulation scores around boundaries were calculated using the deepTools computeMatrix function (https://github.com/deeptools/deepTools).

*Chromatin loops.* The chromatin loops were identified using the Juicer HiCCUPS algorithm with default parameters and the chromosight (https://github.com/koszullab/chromosight) package at 10-kb and 5-kb resolution. Because chromosight has much higher detection sensitivity than HiCCUPS, the chromatin loops called by chromosight were used for analyses of topoisomerase-inhibition Micro-C datasets. Chromatin loops between techniques and conditions were overlapped using the bedtools pairtopair function with "-slop 20000". We also calculated chromatin loops on merged single-cell data at 10-kb resolution using the SnapHiC package (https://github.com/HuMingLab/SnapHiC), a software tailored for single-cell Hi-C data. Pileup analyses of chromatin loops were performed using the coolpuppy package (https://github.com/open2c/coolpuppy).

*Chromatin stripes*. We called chromatin stripes using two different algorithms, Stripenn (https://github.com/ysora/stripenn) and StripeCaller (https://github.com/XiaoTaoWang/StripeCaller) at 5-kb resolution. For StripeCaller, the chromatin stripes were extended to the main diagonal of the contact matrix, and redundant calls anchored at the same locus were merged into a single stripe with maximum length. The comparisons of chromatin stripes between different techniques and conditions were then performed based on the stripe anchor and orientation.

*Pileup analysis of gene structure*. Rescaled pileup analyses for genes were performed using coolpuppy with "--flip-negative-strand –rescale –local". We selected genes with gene length ≥ 50kb. For gene analyses based on transcription level, expressed genes with FPKM ≥ 0.5 were classified into high (80-100%), medium (60-80%), and low (0-60%) group based on the RNAPII ChIP signals of the whole gene body.

*Nucleosome occupancy*. The nucleosome occupancy signals were extracted from the mapped reads. Briefly, PCR duplicates were filtered out from raw SAM files using the samblaster package with "--ignoreUnmated -r". The DANPOS2 package (https://sites.google.com/site/danposdoc/) was then used to calculate the nucleosome occupancy signals with "dpos -a 5 –count 1000000". The average signals of nucleosome occupancy around genomic elements of interest were then calculated with computeMatrix and normalized by setting its mean to 1.

***Analysis of single-cell Micro-C data***

*Generation of contact maps.* Single-cell contact maps were generated from raw sequencing data as we previously described (*19*), using the hickit and dip-c packages. Cells with >45% interchromosomal contacts were excluded (15 out of 355 cellst).

*Generation of 3D genomes.* Genomic regions with chromosome abnormalities were excluded for contacts and removed from the 3D genomes afterwards. Single-cell 3D structures were generated as we previously described, with the hickit package (with parameters "-M" and "Sr1m -c1 -r10m -c2 -b4m -b1m -b200k -D5 -b50k -D5 -b20k -D5 -b10k -D5 -b5k"). We generated 5 replicate structures for each cell with different random seeds (1-5). Repetitive regions were also removed from the 3D structure with "dip-c clean3". Similar to our previous studies, each 20-kb particle represents a radius of ~100 nm (~85 nm for each 10-kb particle and ~68 nm for each 5-kb particle).

*Chromosome territory, A/B compartment, chromosome intermingling and chromatin loops in 3D structures*. For each cell, a root mean squared (r.m.s.) r.m.s.d. (across all particles) was calculated with "dip-c align" at 20-kb, 10-kb and 5-kb resolution. Structures with root mean squared r.m.s.d. ≤ 2 particle radii were retained. Each cell was converted to an mmCIF file with "dip-c color" ("dip-c color -n hg38.chr.txt", "dip-c color -d3", and "dip-c color -c hg38.cpg.${resolution/1000}k.txt") and "dip-c vis" ("dip-c vis -c") for visualization in PyMol (fig. S4 and S5).

*Distance matrix analysis.* For single-cell distance matrix analysis, the region of interest was extracted from the full 3D genome file with "dip-c reg3". For each region of interest, we calculated root mean squared r.m.s.d. (across all particles) for each cell, and structures with root mean squared r.m.s.d ≤ 1.5 were retained for further analysis. This approach allows us to use more cells and increase the sample size. Each representative structure was then converted to an mmCIF file with "dip-c color" ("dip-c color -l hg38.chr.len") and "dip-c vis" for visualization in PyMol.

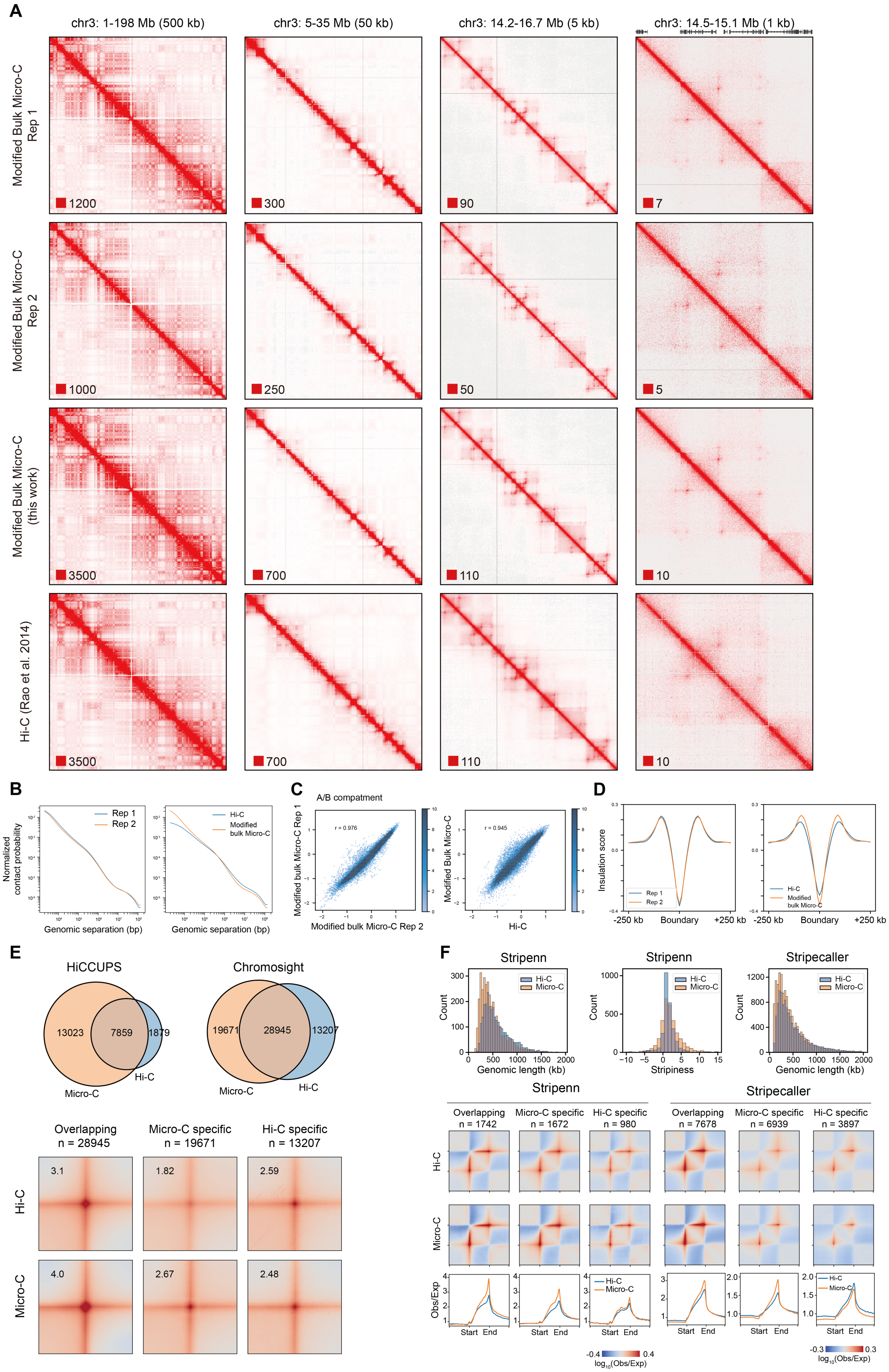
A
chr3: 1-198 Mb (500 kb)
chr3: 5-35 Mb (50 kb)
chr3: 14.2-16.7 Mb (5 kb)
chr3: 14.5-15.1 Mb (1 kb)
Modified Bulk Micro-C Rep 1
1200
300
90
7
Modified Bulk Micro-C Rep 2
1000
250
50
5
Modified Bulk Micro-C (this work)
3500
700
110
10
Hi-C (Rao et al. 2014)
3500
700
110
10
B
Normalized contact probability
Rep 1
Rep 2
Hi-C
Modified bulk Micro-C
Genomic separation (bp)
Genomic separation (bp)
C
A/B compatment
r = 0.976
r = 0.945
Modified bulk Micro-C Rep 1
Modified bulk Micro-C Rep 2
Modified Bulk Micro-C
Hi-C
D
Insulation score
Rep 1
Rep 2
Hi-C
Modified bulk Micro-C
-250 kb
Boundary
+250 kb
-250 kb
Boundary
+250 kb
E
HiCCUPS
13023
7859
1879
Micro-C
Hi-C
Chromosight
19671
28945
13207
Micro-C
Hi-C
Overlapping n = 28945
Micro-C specific n = 19671
Hi-C specific n = 13207
Hi-C
3.1
1.82
2.59
Micro-C
4.0
2.67
2.48
F
Stripenn
Stripenn
Stripecaller
Hi-C
Micro-C
Count
Genomic length (kb)
Stripiness
Genomic length (kb)
Stripenn
Stripecaller
Overlapping n = 1742
Micro-C specific n = 1672
Hi-C specific n = 980
Overlapping n = 7678
Micro-C specific n = 6939
Hi-C specific n = 3897
Hi-C
Micro-C
Obs/Exp
Start
End
-0.4
0.4
log10(Obs/Exp)
-0.3
0.3
log10(Obs/Exp)

**Fig. S1. Quality control of modified bulk Micro-C and comparison with Hi-C.** (**A**) Contact map of two Micro-C replicates and pooled bulk Micro-C and Hi-C at different resolution. (**B**) Interaction frequency as a function of genomic distance between interaction fragments for two Micro-C replicates, bulk Micro-C and Hi-C. (**C**) Compartment are highly consistent between two Micro-C replicates, Micro-C and Hi-C. Scatter plot of genome-wide eigenvector scores. (**D**) Global average insulation scores of TADs boundary for two Micro-C replicates, Micro-C and Hi-C. (**E**) Bulk Micro-C show high sensitivity in chromatin loop detection. Venn diagram of chromatin loops detected between bulk Micro-C and Hi-C (upper panel), two loop caller were used, HiCCUPS and Chromsight. Pile-up analysis of loop enrichment (bottom panel). (**F**) Bulk Micro-C detects more stripe structures than Hi-C, two stripe caller were used for benchmark, Stripenn and Stripecaller. Pile-up analysis of stripe enrichment and quantification are plotted.

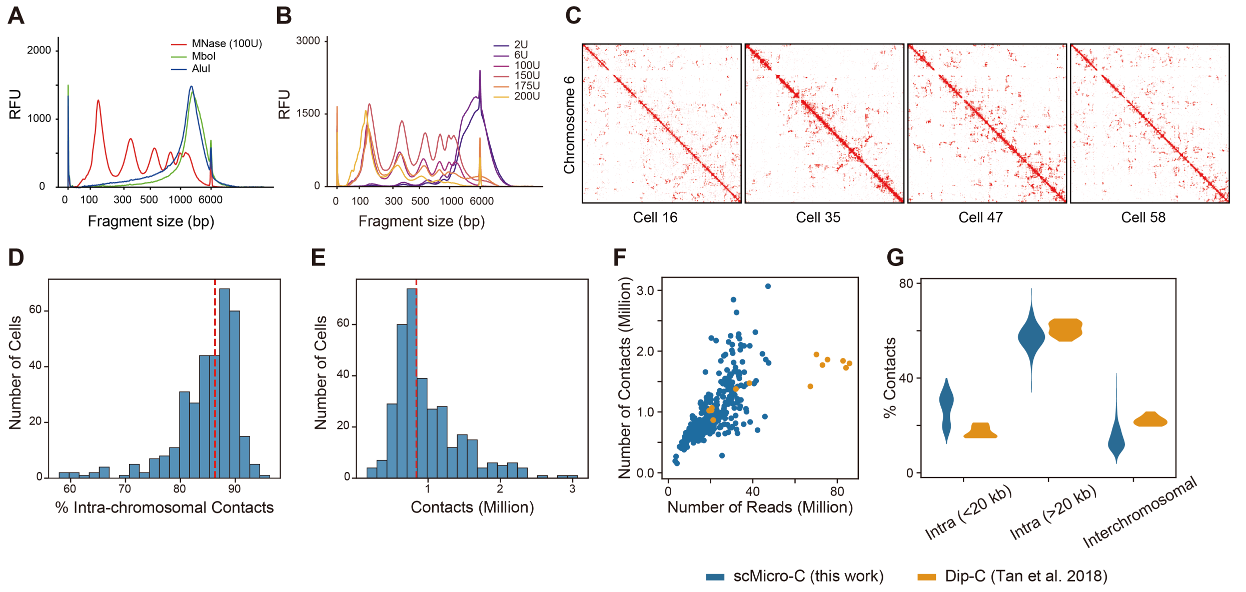


**Fig. S2. Validation of the single-cell Micro-C method.** (**A**) Fragment length distribution after digestion of two restriction enzymes or MNase (data from capillary electrophoresis). (**B**) Fragment length distribution of MNase titration on 1 million GM12878 cells. (**C**) Single-cell contact maps of GM12878 cells from chromosome 6 at 500-kb resolution. (**D**) Histogram of the intrachromosomal contact ratio. (**E**) Histogram of the numbers of contacts per cell. (**F**) The numbers of contacts per cell against the numbers of raw sequencing read pairs for scMicro-C and Dip-C. (**G**) Ther percentage of short-range (≤20 kb), long-range contacts (≥20 kb) and interchromosomal contacts for scMicro-C and Dip-C.

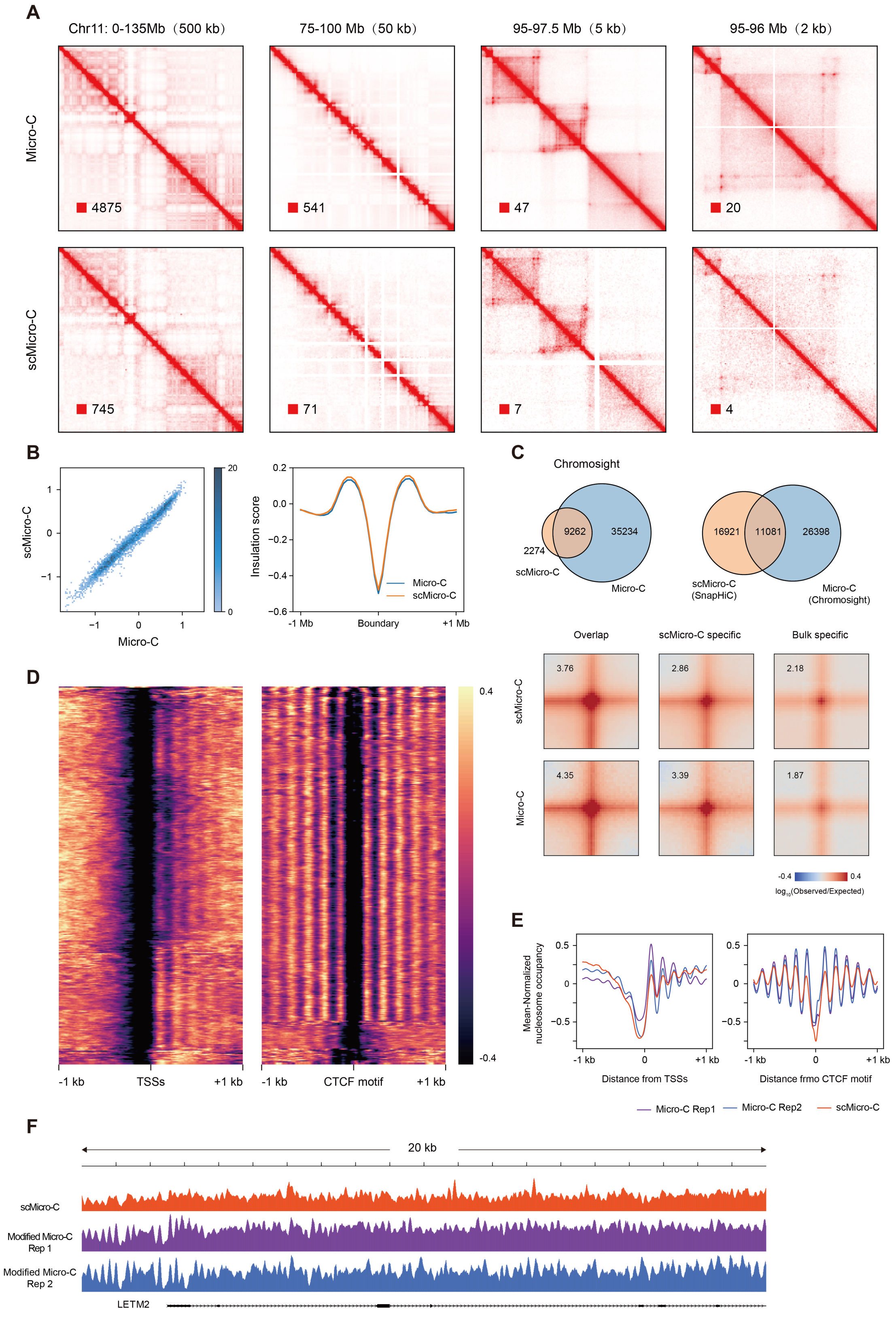
A
Chr11: 0-135Mb（500 kb）
75-100 Mb（50 kb）
95-97.5 Mb（5 kb）
95-96 Mb（2 kb）
Micro-C
scMicro-C
4875
541
47
20
745
71
7
4
B
scMicro-C
Micro-C
Insulation score
-1 Mb
Boundary
+1 Mb
Micro-C
scMicro-C
C
Chromosight
2274
9262
35234
scMicro-C
Micro-C
16921
11081
26398
scMicro-C (SnapHiC)
Micro-C (Chromosight)
Overlap
scMicro-C specific
Bulk specific
3.76
2.86
2.18
4.35
3.39
1.87
log10(Observed/Expected)
D
-1 kb
TSSs
+1 kb
-1 kb
CTCF motif
+1 kb
E
Mean-Normalized nucleosome occupancy
Distance from TSSs
Distance frmo CTCF motif
Micro-C Rep1
Micro-C Rep2
scMicro-C
F
20 kb
scMicro-C
Modified Micro-C Rep 1
Modified Micro-C Rep 2
LETM2

**Fig. S3. scMicro-C retains characteristics of Micro-C data.** (**A**) Contact map of ensemble scMicro-C and bulk Micro-C at different resolution. (**B**) Eigenvector values are highly correlated between scMicro-C and bulk Micro-C genome-wide, scatter plot shows the first eigenvector values at 100 kb resolution (left). TADs are highly consistent between scMicro-C and bulk Micro-C genome-wide, scatter plot of insulation scores of TADs boundary at 50 kb resolution (right). (**C**) Venn diagram of chromatin loops detected with ensemble scMicro-C and bulk Micro-C, scMicro-C data also use SnapHi-C algorithm (upper panel). Pile-up analysis of loop enrichment. (**D**) Nucleosome occupancy around active transcription start sites (TSSs) and CTCF-binding sites among single cells. (**E**) Mean nucleosome occupancy of ensemble scMicro-C and modified bulk Micro-C around TSS and CTCF-binding sites. (**F**) Nucleosome occupancy tracks for ensemble scMicro-C (top) and two replicates of modified bulk Micro-C (bottom).

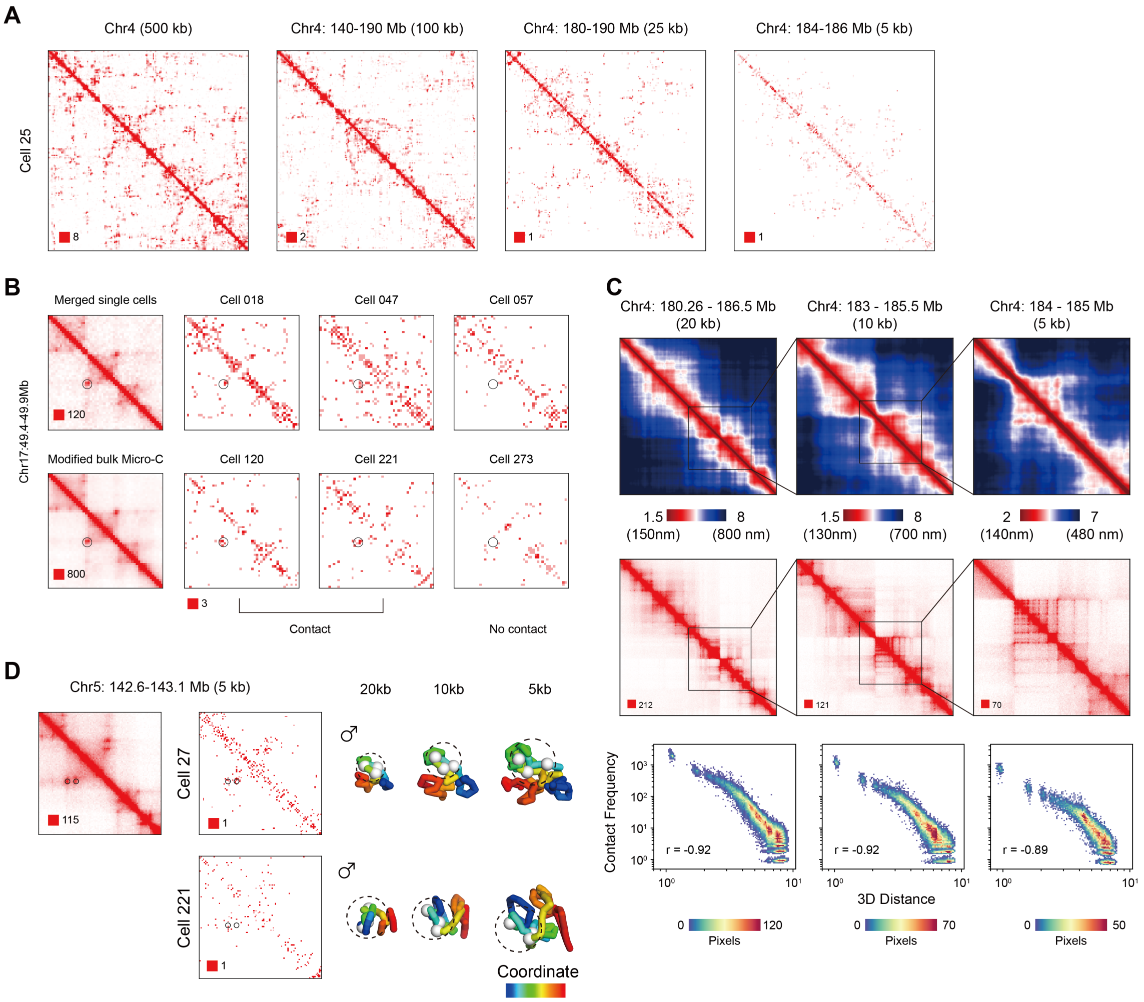


**Fig. S4. Validation of high-resolution 3D genome structures.** (**A**) Contact map of a single cell at different resolution. (**B**)Contact map of a loop structure, bulk Micro-C, ensemble scMicro-C and single cells. (**C**)The mean distance matrices (upper) agree well with the bulk contact maps (middle) for the corresponding regions at different resolutions, scatter plot of contact probability and mean spatial distance at different resolutions (bottom). (**D**) 3D single-cell structures of a loop region at 20 kb, 10 kb and 5 kb resolution, single-cell contact maps are shown.

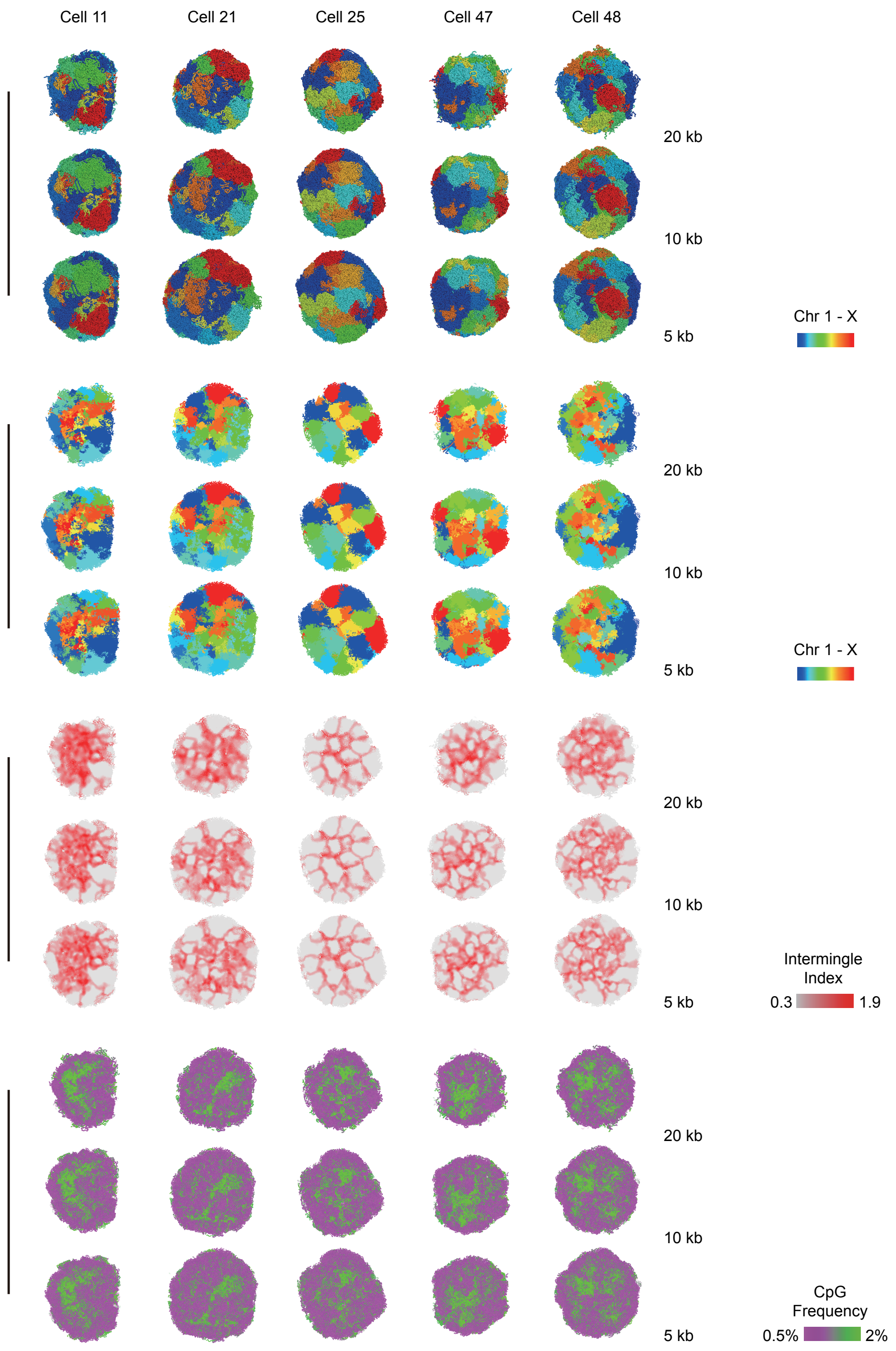


**Fig. S5. High-resolution 3D genome structures of GM12878 cells.** Whole genome structures of 5 cells at 20 kb, 10 kb and 5 kb resolution, with particles colored according to chromosomes, multi-chromosome intermingle index or CpG frequency (from top to bottom).

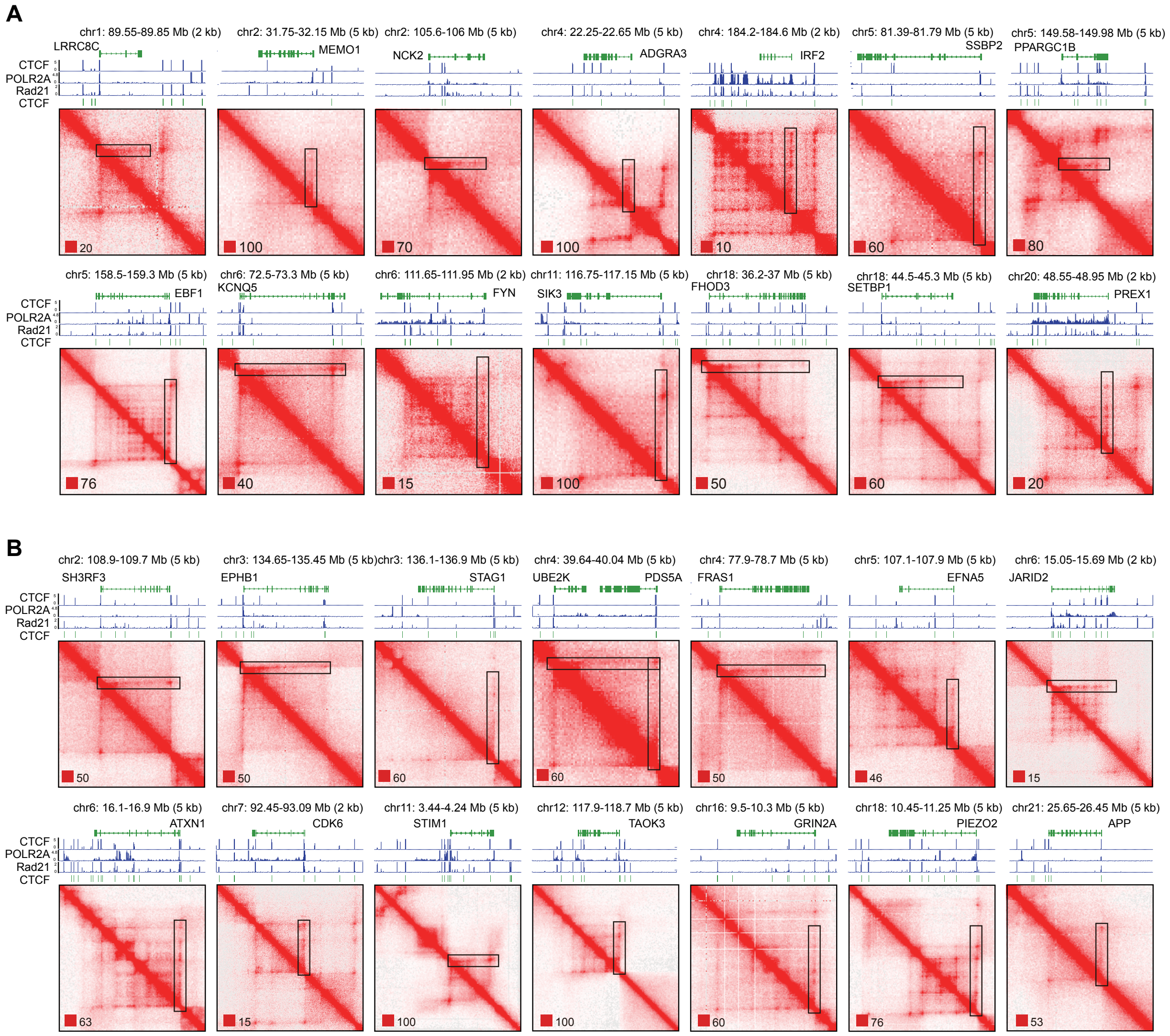


**Fig. S6. Identification of transcription elongation loops.** (**A**) Contact maps of TELs without CTCF binding at TSS region. (**B**) Contact maps of TELs with CTCF binding at TSS region.

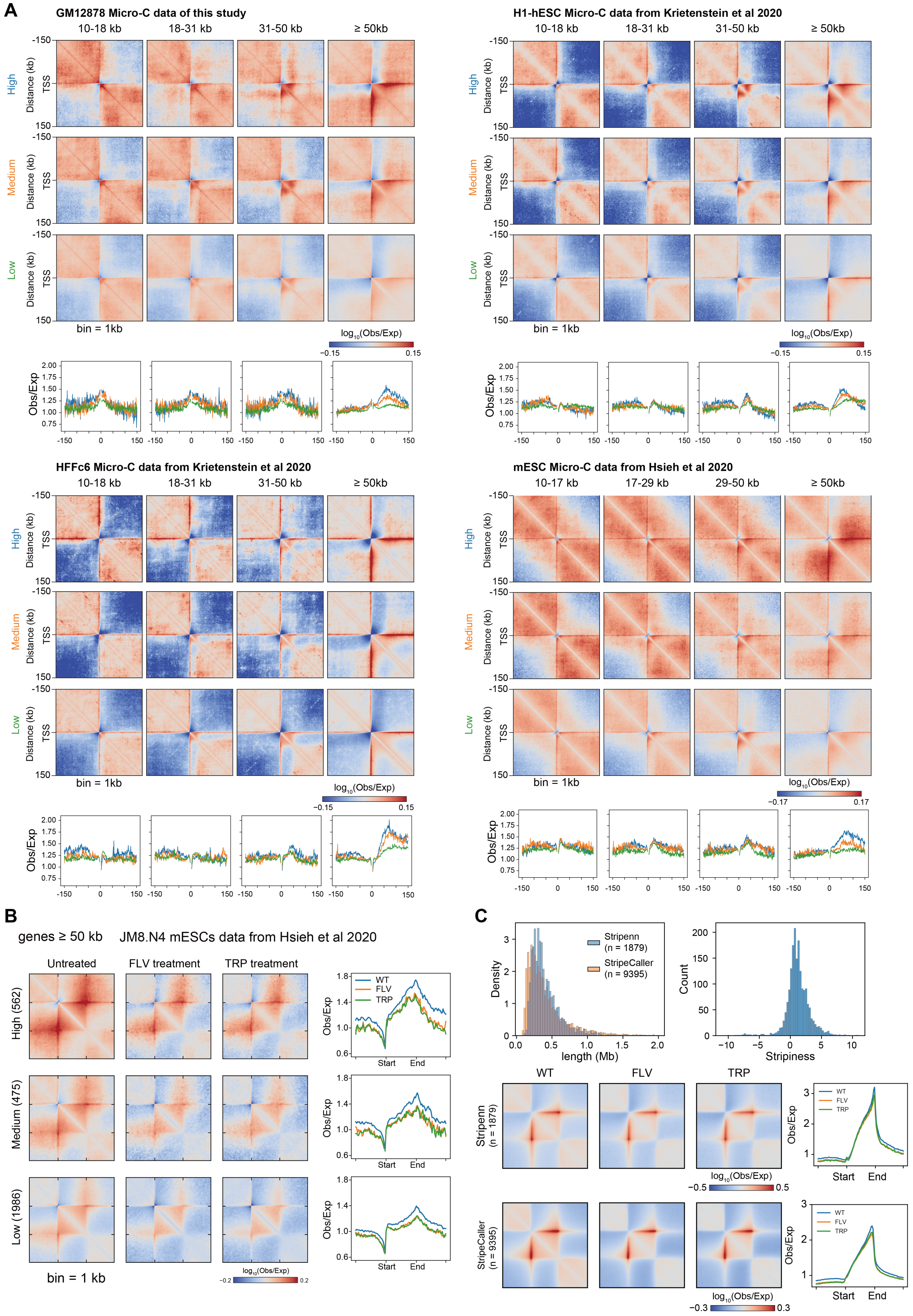

A
GM12878 Micro-C data of this study
10-18 kb
18-31 kb
31-50 kb
≥ 50kb
H1-hESC Micro-C data from Krietenstein et al 2020
HFFc6 Micro-C data from Krietenstein et al 2020
mESC Micro-C data from Hsieh et al 2020
10-17 kb
17-29 kb
29-50 kb
High
Medium
Low
TSS
Distance (kb)
bin = 1kb
log10(Obs/Exp)
Obs/Exp
B
genes ≥ 50 kb
JM8.N4 mESCs data from Hsieh et al 2020
Untreated
FLV treatment
TRP treatment
High (562)
Medium (475)
Low (1986)
bin = 1 kb
WT
FLV
TRP
Start
End
C
Stripenn (n = 1879)
StripeCaller (n = 9395)
Density
length (Mb)
Count
Stripiness

**Fig. S7. TELs are dependent on gene length and transcription activity.** (**A**) Pile-up analysis of four Micro-C datasets (*16, 17*), genes are grouped into different length bins and transcriptional activity. (**B**) Acute transcription inhibition disrupts TELs. Rescaled pile-up analysis of acute transcription inhibition by triptolide (TRP) or flavopiridol (FLV), generated with Micro-C of mESCs cell line (*16*) at 1 kb resolution. genes are sorted by transcriptional activity into high (n = 562), medium (n = 475) and low (n = 1986). Only long genes (≥ 50 kb) are shown (left). Quantification of TSS interaction enrichment (right). (**C**) Architectural stripes are largely unaffected upon transcription inhibition. Stripe scores and length distribution of stripes called in untreated mESCs Micro-C dataset (top). Pile-up analysis of interaction enrichment at stripes of untreated and transcription inhibition datasets using stripes called with Stripenn(middle) and StripeCaller (bottom). Quantification of interaction enrichment at stripe region are shown on right respectively.

A

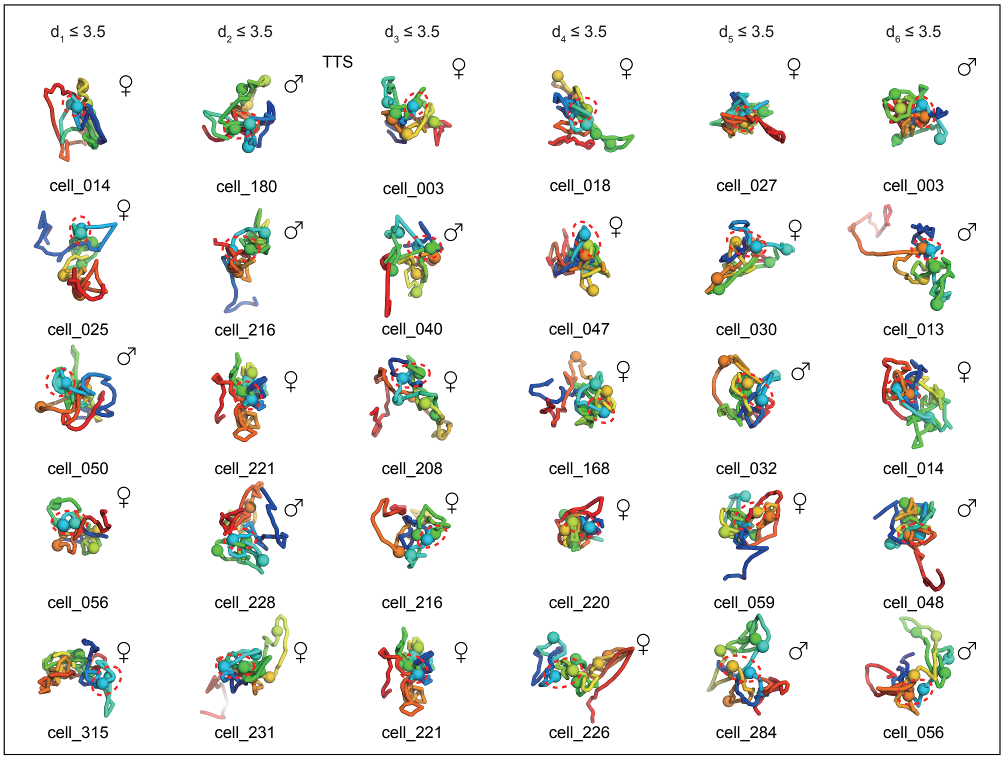
$d_1 \leq 3.5$
$d_2 \leq 3.5$
$d_3 \leq 3.5$
$d_4 \leq 3.5$
$d_5 \leq 3.5$
$d_6 \leq 3.5$
TTS
cell_014
cell_180
cell_003
cell_018
cell_027
cell_003
cell_025
cell_216
cell_040
cell_047
cell_030
cell_013
cell_050
cell_221
cell_208
cell_168
cell_032
cell_014
cell_056
cell_228
cell_216
cell_220
cell_059
cell_048
cell_315
cell_231
cell_221
cell_226
cell_284
cell_056

B

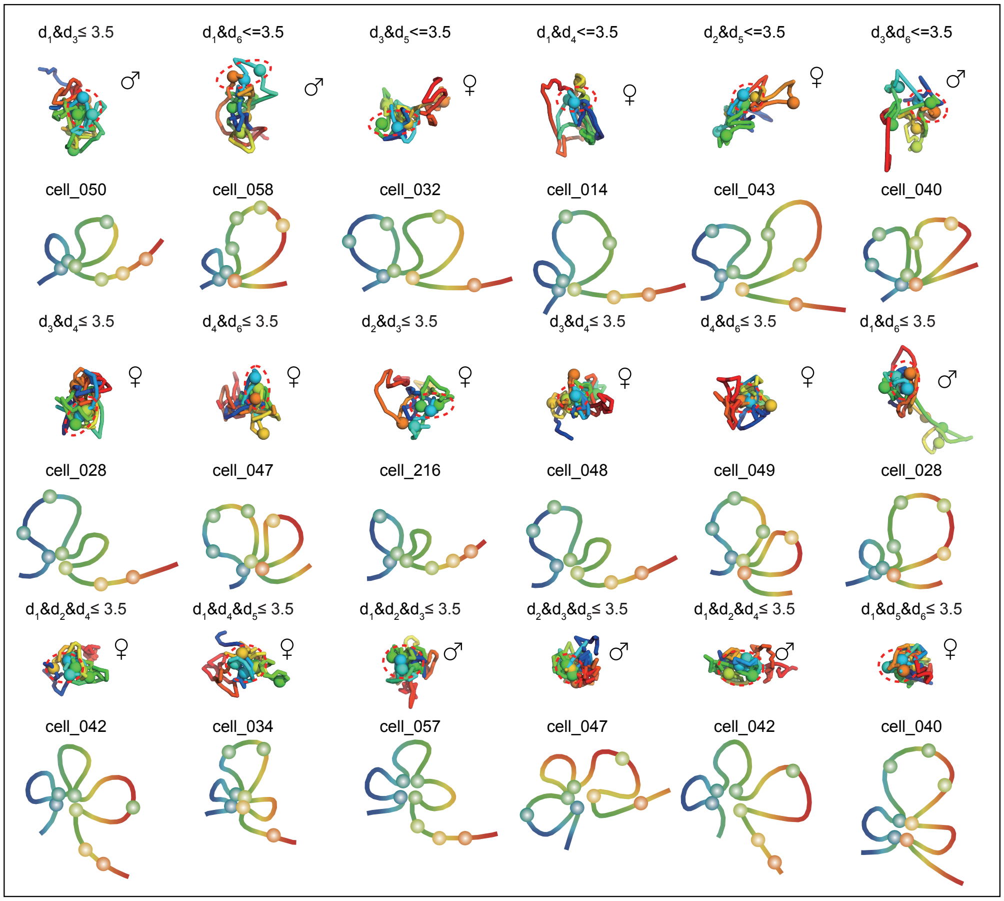
$d_1 \& d_3 \leq 3.5$
$d_1 \& d_6 <= 3.5$
$d_3 \& d_5 <= 3.5$
$d_1 \& d_4 <= 3.5$
$d_2 \& d_5 <= 3.5$
$d_3 \& d_6 <= 3.5$
cell_050
cell_058
cell_032
cell_014
cell_043
cell_040
$d_3 \& d_4 \leq 3.5$
$d_4 \& d_6 \leq 3.5$
$d_2 \& d_3 \leq 3.5$
$d_3 \& d_4 \leq 3.5$
$d_4 \& d_6 \leq 3.5$
$d_1 \& d_6 \leq 3.5$
cell_028
cell_047
cell_216
cell_048
cell_049
cell_028
$d_1 \& d_2 \& d_4 \leq 3.5$
$d_1 \& d_4 \& d_5 \leq 3.5$
$d_1 \& d_2 \& d_3 \leq 3.5$
$d_2 \& d_3 \& d_5 \leq 3.5$
$d_1 \& d_2 \& d_4 \leq 3.5$
$d_1 \& d_5 \& d_6 \leq 3.5$
cell_042
cell_034
cell_057
cell_047
cell_042
cell_040

**Fig. S8. Recapitulating TELs folding with kilobase-resolution single-cell 3D structures.** (**A**) Representative single-cell structures following gradual loop extension process. (**B**)Single-cell structures with multi-way interactions, corresponding schematic depictions are show.

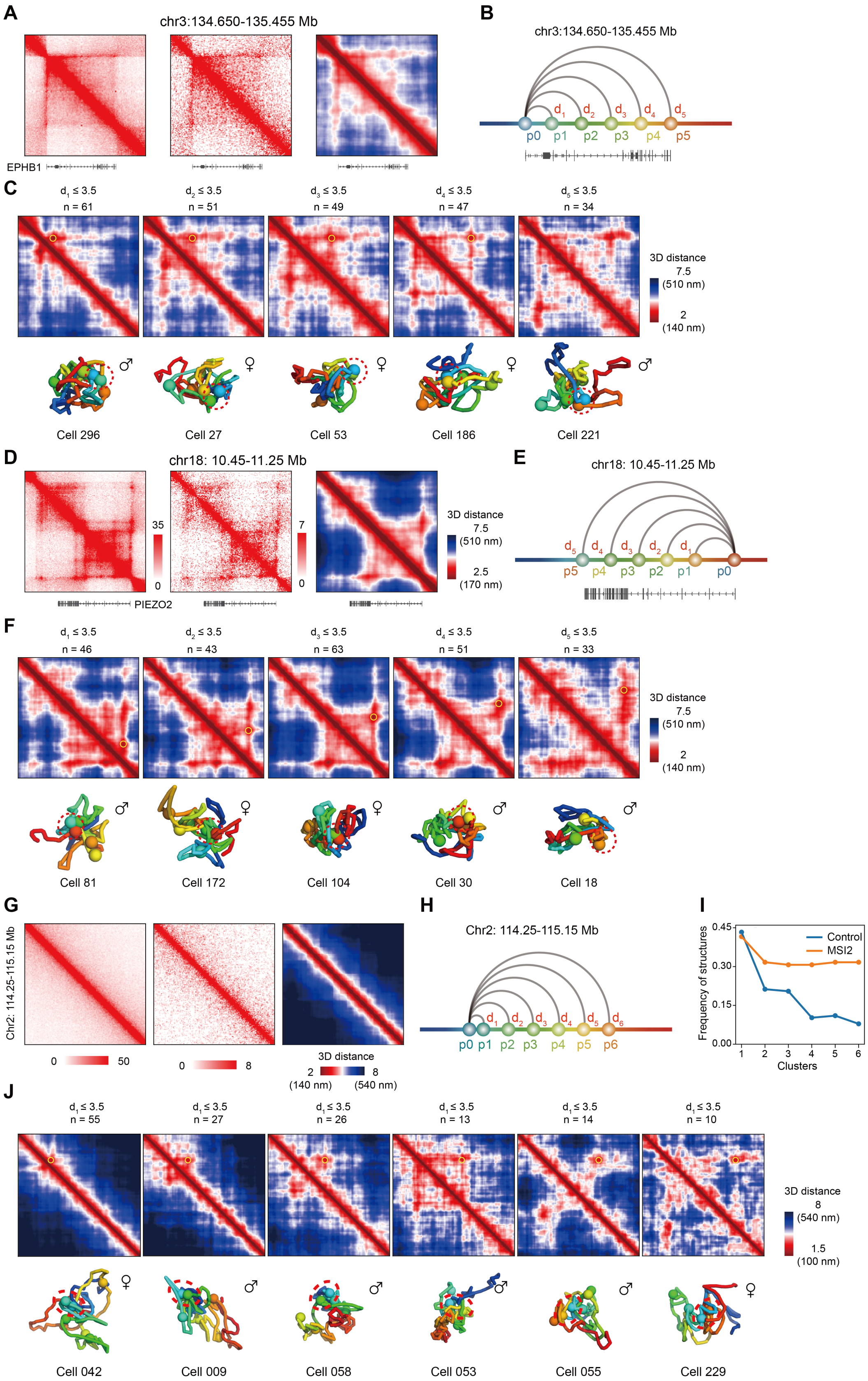
A
chr3:134.650-135.455 Mb
EPHB1
B
chr3:134.650-135.455 Mb
d1 d2 d3 d4 d5
p0 p1 p2 p3 p4 p5
C
d1 ≤ 3.5 n = 61
d2 ≤ 3.5 n = 51
d3 ≤ 3.5 n = 49
d4 ≤ 3.5 n = 47
d5 ≤ 3.5 n = 34
3D distance
7.5 (510 nm)
2 (140 nm)
Cell 296
Cell 27
Cell 53
Cell 186
Cell 221
D
chr18: 10.45-11.25 Mb
35
0
7
0
PIEZO2
3D distance
7.5 (510 nm)
2.5 (170 nm)
E
chr18: 10.45-11.25 Mb
d5 d4 d3 d2 d1
p5 p4 p3 p2 p1 p0
F
d1 ≤ 3.5 n = 46
d2 ≤ 3.5 n = 43
d3 ≤ 3.5 n = 63
d4 ≤ 3.5 n = 51
d5 ≤ 3.5 n = 33
3D distance
7.5 (510 nm)
2 (140 nm)
Cell 81
Cell 172
Cell 104
Cell 30
Cell 18
G
Chr2: 114.25-115.15 Mb
0 50
0 8
3D distance
2 (140 nm)
8 (540 nm)
H
Chr2: 114.25-115.15 Mb
d1 d2 d3 d4 d5 d6
p0 p1 p2 p3 p4 p5 p6
I
Frequency of structures
0.45
0.30
0.15
0.00
Control
MSI2
1 2 3 4 5 6
Clusters
J
d1 ≤ 3.5 n = 55
d1 ≤ 3.5 n = 27
d1 ≤ 3.5 n = 26
d1 ≤ 3.5 n = 13
d1 ≤ 3.5 n = 14
d1 ≤ 3.5 n = 10
3D distance
8 (540 nm)
1.5 (100 nm)
Cell 042
Cell 009
Cell 058
Cell 053
Cell 055
Cell 229

**Fig. S9. Single-cell 3D structures of stripe and non-stripe regions.** (**A**) Contact maps of bulk Micro-C (left) and ensemble scMicro-C (middle) at around EPHB1 gene (chr3: 134.650-135.455 Mb) at 5 kb resolution. And average distance matrix of all avaible single-cell 3D structures (right). Gene track is shown on the bottom. (**B**) Schematic of selected genomic loci for 3D structure distance analysis (p0 to p5). (**C**) Average distance matrix of structures belonging to five clusters with 3D distance less than 3.5 particle radii (~240 nm) to reference point (p0) (top). Representative single-cell 3D structures are shown (bottom). (**D**-**F**) The same as (**A**-**C**), for PIEZO2 gene. (**G**) Contact map of bulk Micro-C (left) and ensemble scMicro-C (middle) at chr2: 11.435-11.515 Mb at 5 kb resolution. Average distance matrix of all avaible single-cell 3D structures (right). (**H**) Schematic of genomic loci selection for 3D structure distance analysis (p0-p6). (**I**) Percentage of structures with distance ≤ 3.5 particle radii for MSI2 gene and control region. (**J**)Average distance matrix of six clusters based on supervised clustering according to distance to p0 (top). Representative single-cell 3D structures (bottom).

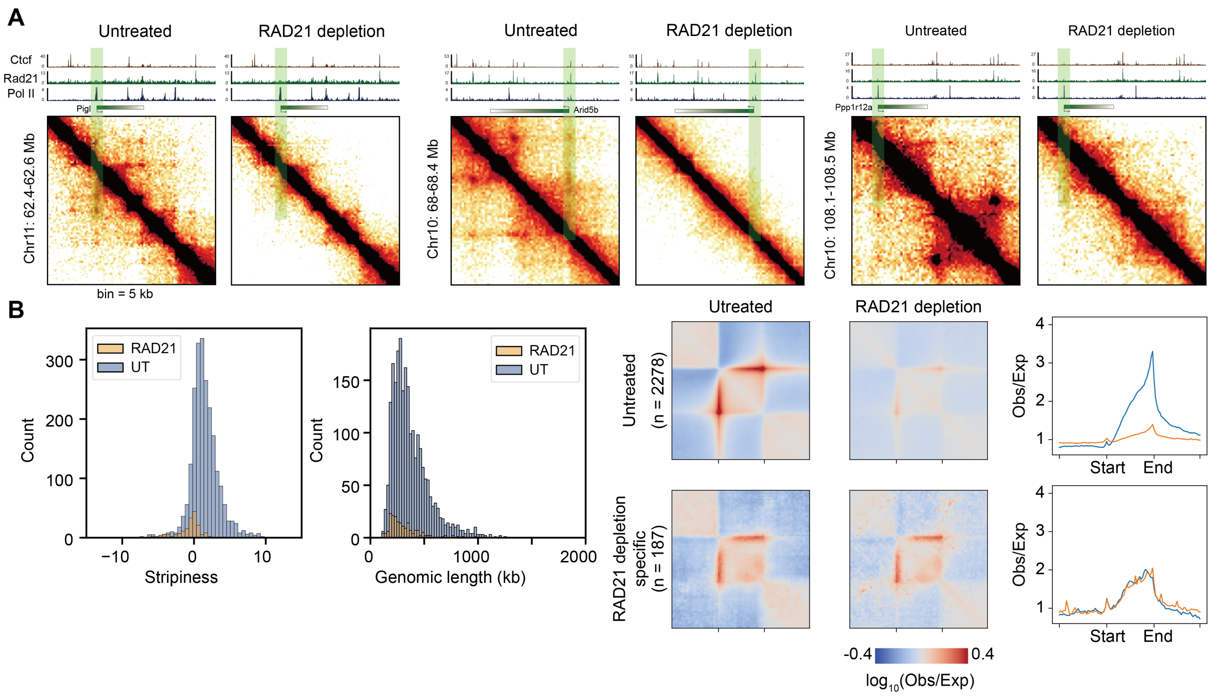


**Fig. S10. TELs and architectural stripes are both dependent on cohesin.** (**A**) Contact map of 3 TELs of untreated and RAD21 depletion, data from Hsieh et al 2022 (*13*). (**B**) RAD21 depletion abolishes almost all architectural stripes. Stripe scores and length distribution of stripes called in untreated, RAD21 depletion datasets (left). Pile-up analysis of interactions at stripes of untreated, RAD21 depletion datasets (middle). Quantification of interaction enrichment at stripe region (right).

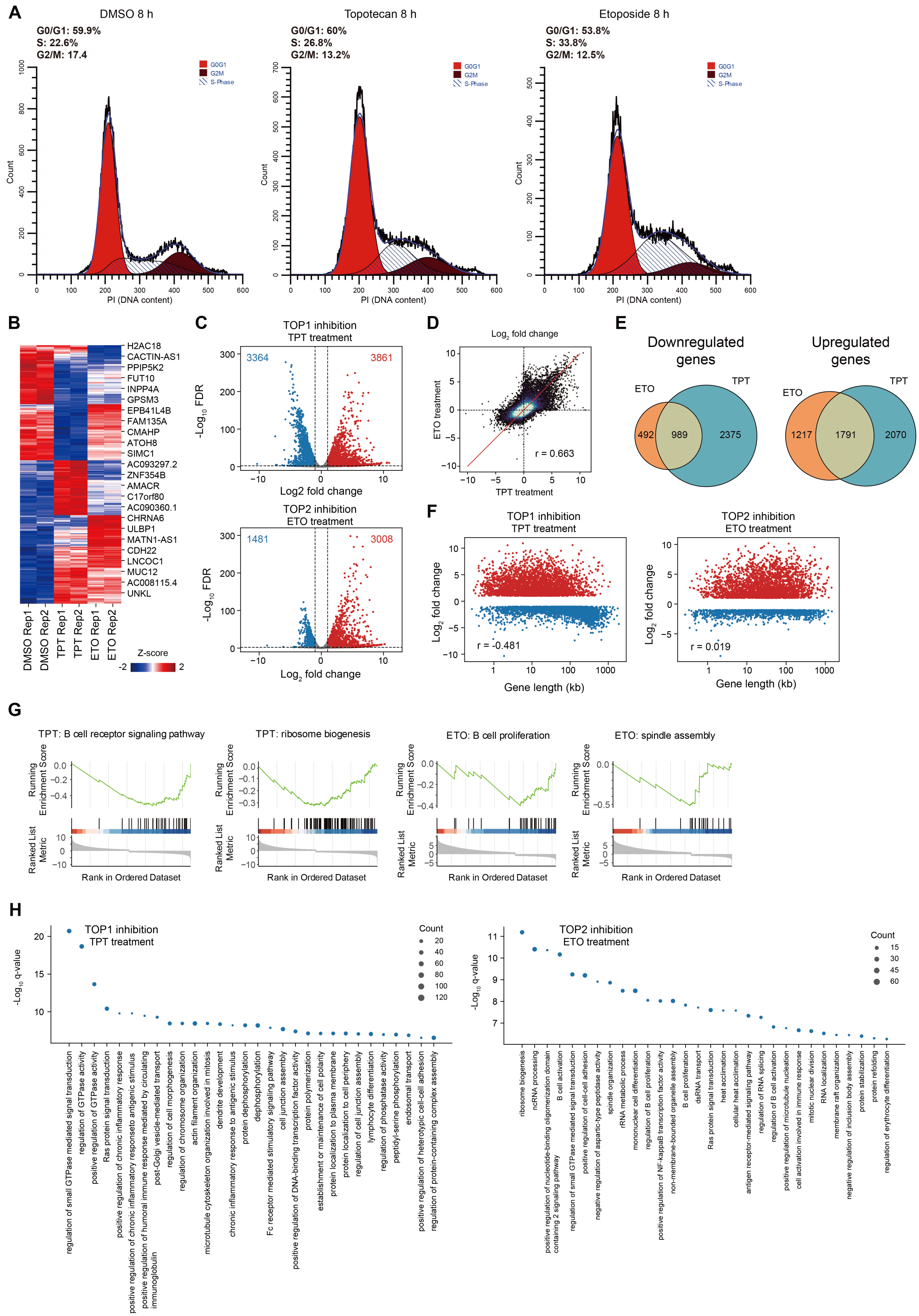
A
DMSO 8 h
G0/G1: 59.9%
S: 22.6%
G2/M: 17.4
Topotecan 8 h
G0/G1: 60%
S: 26.8%
G2/M: 13.2%
Etoposide 8 h
G0/G1: 53.8%
S: 33.8%
G2/M: 12.5%
G0G1
G2M
S-Phase
Count
PI (DNA content)
B
H2AC18
CACTIN-AS1
PPIP5K2
FUT10
INPP4A
GPSM3
EPB41L4B
FAM135A
CMAHP
ATOH8
SIMC1
AC093297.2
ZNF354B
AMACR
C17orf80
AC090360.1
CHRNA6
ULBP1
MATN1-AS1
CDH22
LNCOC1
MUC12
AC008115.4
UNKL
DMSO Rep1
DMSO Rep2
TPT Rep1
TPT Rep2
ETO Rep1
ETO Rep2
Z-score
C
TOP1 inhibition
TPT treatment
3364
3861
-Log10 FDR
Log2 fold change
TOP2 inhibition
ETO treatment
1481
3008
D
Log2 fold change
ETO treatment
TPT treatment
r = 0.663
E
Downregulated genes
ETO
TPT
492
989
2375
Upregulated genes
1217
1791
2070
F
TOP1 inhibition
TPT treatment
Log2 fold change
r = -0.481
Gene length (kb)
TOP2 inhibition
ETO treatment
r = 0.019
G
TPT: B cell receptor signaling pathway
TPT: ribosome biogenesis
ETO: B cell proliferation
ETO: spindle assembly
Running Enrichment Score
Ranked List Metric
Rank in Ordered Dataset
H
TOP1 inhibition
TPT treatment
-Log10 q-value
Count
regulation of small GTPase mediated signal transduction
regulation of GTPase activity
positive regulation of GTPase activity
Ras protein signal transduction
positive regulation of chronic inflammatory response
positive regulation of chronic inflammatory responseto antigenic stimulus
positive regulation of humoral immune response mediated by circulating immunoglobulin
post-Golgi vesicle-mediated transport
regulation of cell morphogenesis
regulation of chromosome organization
actin filament organization
microtubule cytoskeleton organization involved in mitosis
dendrite development
chronic inflammatory response to antigenic stimulus
protein dephosphorylation
dephosphorylation
Fc receptor mediated stimulatory signaling pathway
cell junction assembly
positive regulation of DNA-binding transcription factor activity
protein polymerization
establishment or maintenance of cell polarity
protein localization to plasma membrane
protein localization to cell periphery
regulation of cell junction assembly
lymphocyte differentiation
regulation of phosphatase activity
peptidyl-serine phosphorylation
endosomal transport
positive regulation of heterotypic cell-cell adhesion
regulation of protein-containing complex assembly
TOP2 inhibition
ETO treatment
ribosome biogenesis
ncRNA processing
positive regulation of nucleotide-binding oligomerization domain containing 2 signaling pathway
B cell activation
regulation of small GTPase mediated signal transduction
positive regulation of cell-cell adhesion
negative regulation of aspartic-type peptidase activity
spindle organization
rRNA metabolic process
mononuclear cell differentiation
regulation of B cell proliferation
positive regulation of NF-kappaB transcription factor activity
non-membrane-bounded organelle assembly
B cell proliferation
dsRNA transport
Ras protein signal transduction
heat acclimation
cellular heat acclimation
antigen receptor-mediated signaling pathway
regulation of RNA splicing
regulation of B cell activation
positive regulation of microtubule nucleation
cell activation involved in immune response
mitotic nuclear division
RNA localization
membrane raft organization
negative regulation of inclusion body assembly
protein stabilization
protein refolding
regulation of erythrocyte differentiation

**Fig. S11. Topoisomerase inhibition downregulates highly expressed long genes involving in housekeeping and cell-type-specific processes.** (**A**)Flow cytometry for cells treated with DMSO, TOP1 inhibitor (TPT) and TOP2 inhibitor (ETO) inhibition stained with propidium iodide (PI) to measure DNA content for cell cycle analysis. (**B**) Expression levels of genes upon DMSO (no inhibition), ETO (TOP2 inhibition) and TPT treatment (TOP1 inhibition). (**C**) Volcano plots of differentially expressed genes after ETO and TPT treatment respectively. (**D**) Scatterplot plot of genes showing fold change between ETO and TPT treatment. TOP2 and TOP1 inhibition have similar effects on gene expression. (**E**) Venn diagram of differentially expressed genes between ETO and TPT treatment. (**F**) Scatterplot showing the fold change versus gene length. (**G**) Gene set enrichment analysis of downregulated genes upon TOP1 and TOP2 inhibition. Shown are housekeeping and cell-type-specific pathways. (**H**) Gene ontology analysis of TOP1 inhibition-downregulated and TOP2 inhibition-downregulated genes.

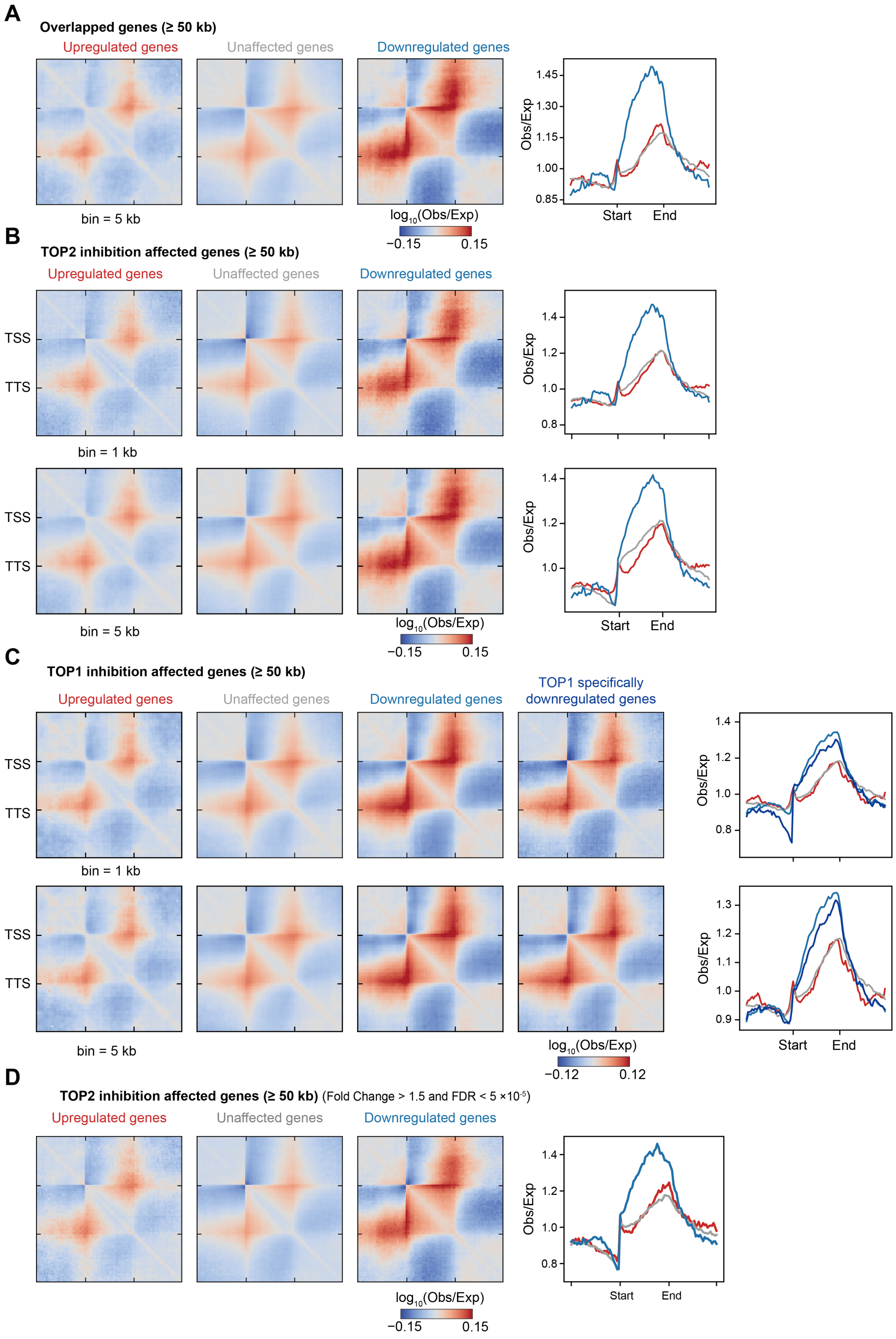


**Fig. S12. Genes downregulated by topoisomerase inhibition have strong TEL signals.** (**A**) Pile-up analysis of overlapped genes between TOP1 and TOP2 inhibition of genes ≥ 50 kb at 5 kb resolution. (**B**) TOP2 inhibition-downregulated genes show strong TEL pattern. Pile-up analysis of genes affected by TOP2 inhibition with cutoff (Fold Change > 2 and FDR < 0.01), upregulated (n = 1052), unaffected (n = 5001) and downregulated (n = 658), at 1 kb resolution (top), and at 5 kb resolution (bottom). Right: lineplot showing the average TEL signal. (**C**) Same analysis as in (**B**) but with genes affected by TOP1 inhibition with cutoff (Fold Change > 2 and FDR < 0.01), upregulated (n = 601), unaffected (n = 3932), downregulated (n = 2169) and TOP1-specifically downregulated genes (n = 1586). TOP1 specifically downregulated genes show weaker TELs signal. (D) Pile-up analysis of genes affected by TOP2 inhibition with cutoff (Fold Change > 1.5 and FDR < 5 x 10^-5). Upregulated (n = 771), unaffected (n = 4807) and downregulated (n = 1133).

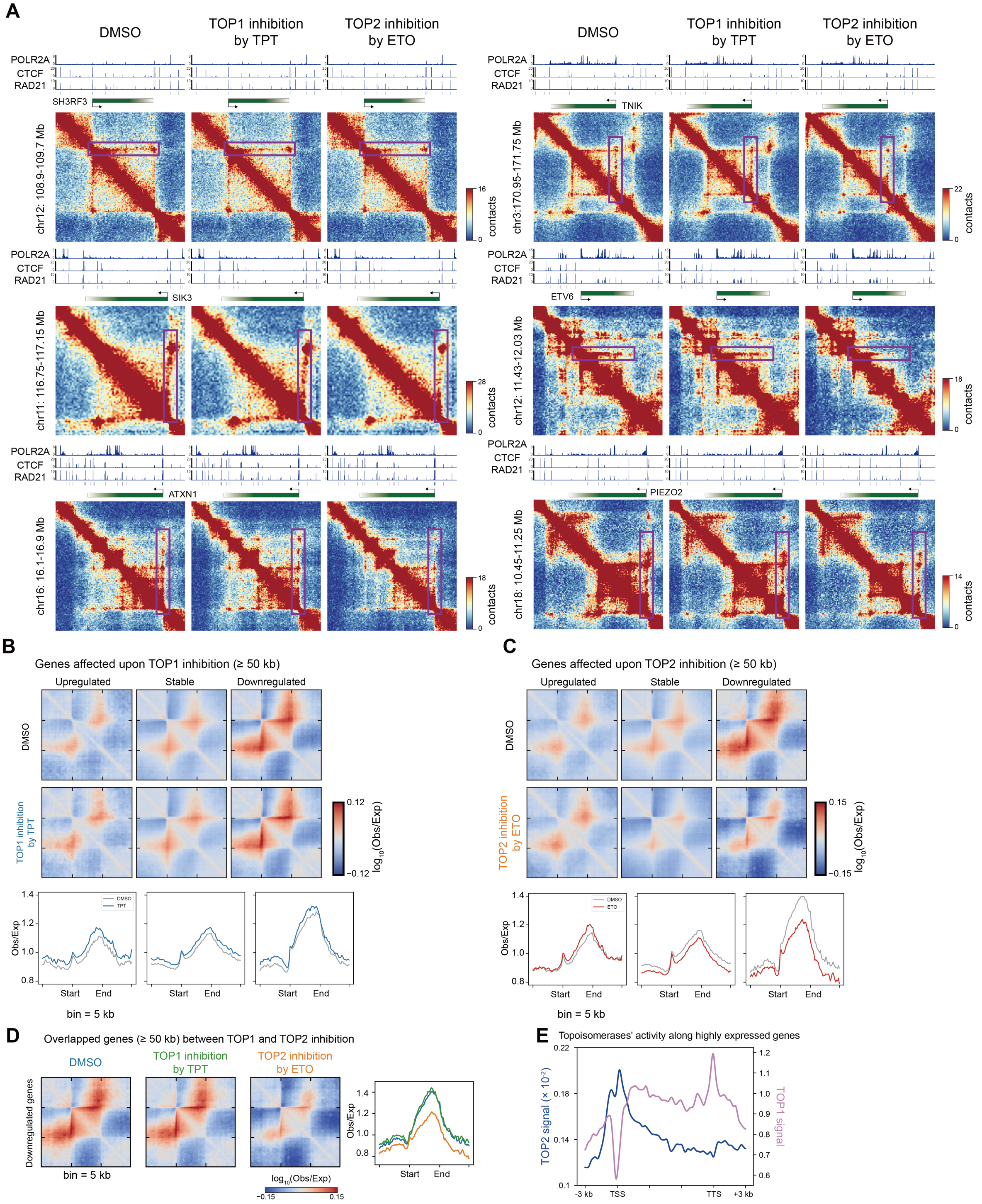


**Fig. S13. TOP1 and TOP2 inhibition differentially affects TELs.** (**A**) Examples of TELs contact matrices of DMSO (no topoisomerase inhibition), ETO (TOP2 inhibition) and TPT treatment (TOP1 inhibition) Micro-C datasets. Vertical boxes denote 3' TELs; Horizontal boxes denote 5' TELs. (**B**) Pile-up analysis of GM12878 Micro-C datasets treated with control (DMSO) and TOP1 inhibitor (TPT), genes are grouped based on RNA-seq of TOP1 inhibition. (**C**) Pile-up analysis of GM12878 Micro-C datasets treated with control (DMSO) and TOP2 inhibitor (TPT), genes are grouped based on RNA-seq of TOP2 inhibition. (**D**) The same as Fig. 3C but at 5 kb resolution. (E) The activity of TOP1 and TOP2 along highly expressed genes. TOP1 activity use TOP1-seq data in HCT116 cells from Baranello et al 2016 (*30*), only highly expressed genes (95-100%) are plotted. TOP2 activity use CC-seq data in human RPE-1 cells from Gittens et al 2019 (*31*), only the top 1/4 highly expressed genes are plotted.

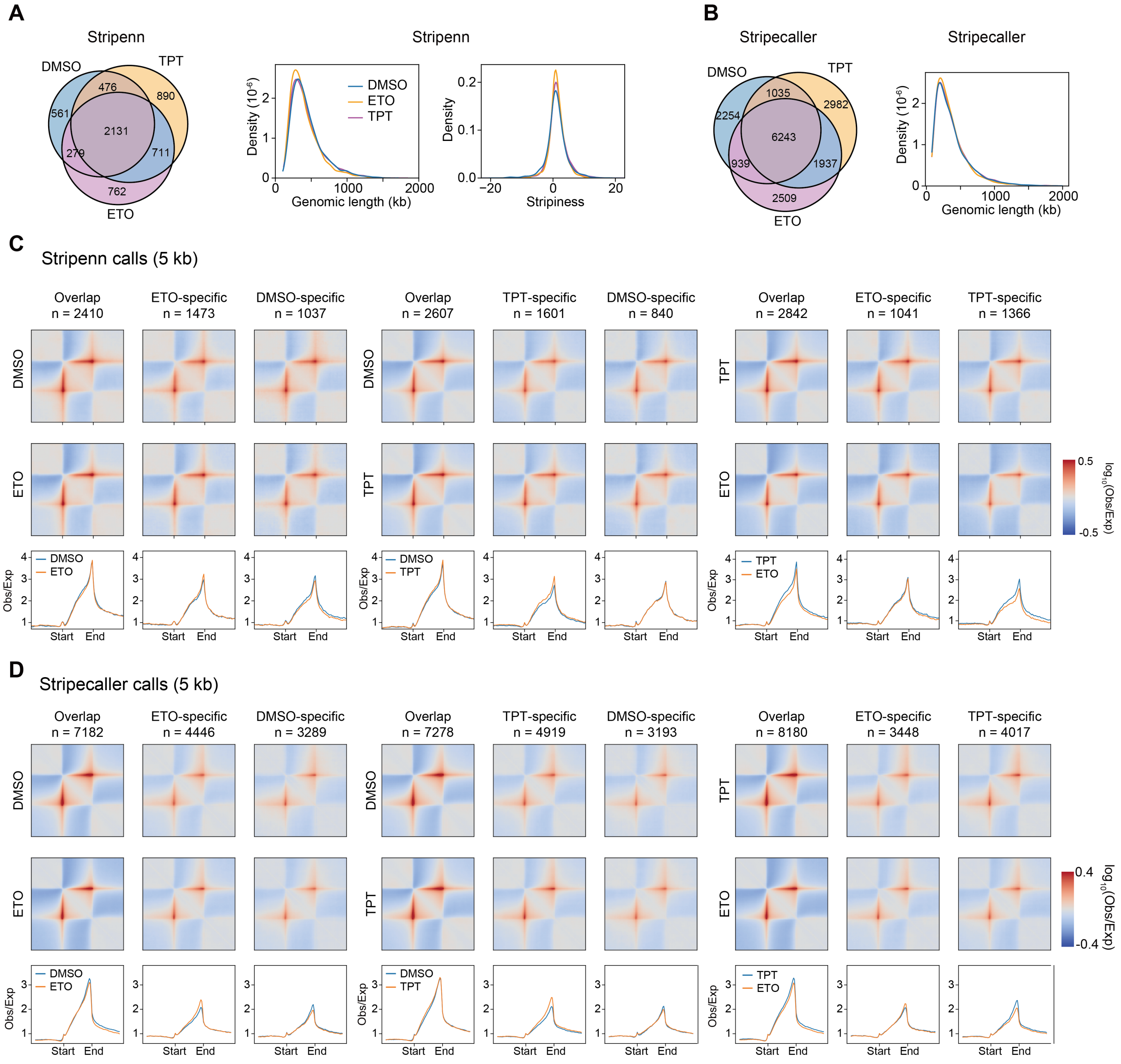


**Fig. S14. Architectural stripes are largely unaffected by topoisomerase inhibition.** (**A**) Chromatin stripes called by Stripenn. Left: overlaps in chromatin stripes between TPT treatment, ETO treatment and DMSO control; middle: length distribution of stripes; right: stripe scores defined by Stripenn. (**B**) Chromatin stripes called by Stripecaller. Left: Overlaps in chromatin stripes between TPT treatment, ETO treatment and DMSO control; right: legth distribution of chromatin stripes. (**C**) Pileup analysis of Stripenn results showing average contact strength of chromatin stripes is largely unaffected after either TOP1 inhibition (TPT treatment) or TOP2 inhibition (ETO treatment). Line plots showing average contact strength of chromatin stripes. (**D**) same analysis as in (**C**) but chromatin stripes called using Stripecaller results.

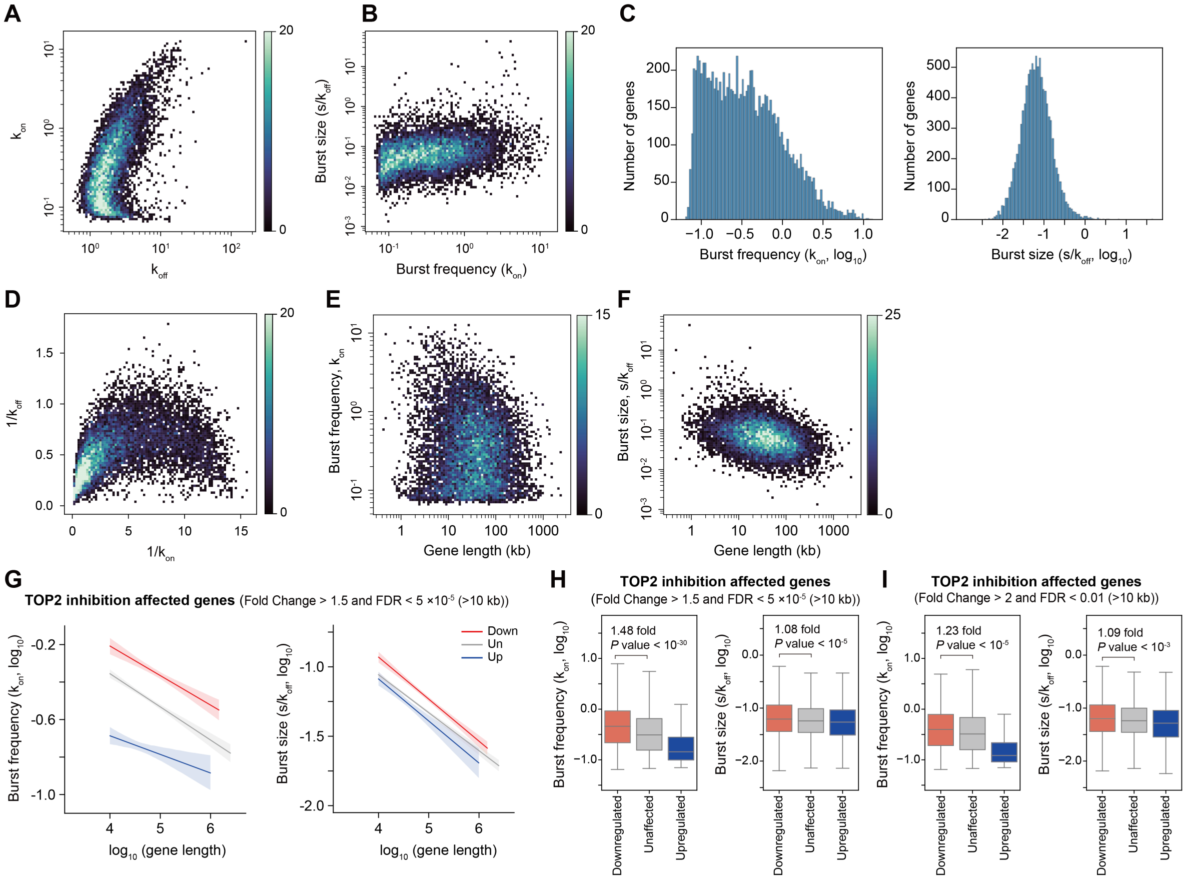


**Fig. S15. TELs ensure higher transcriptional burst frequencies.** (**A**) Scatterplot of transcriptome-wide values of $k_{on}$ and $k_{off}$. (**B**) Scatterplot of transcriptome-wide values of burst frequency and burst size. (**C**) Distribution of transcriptome-wide values of burst frequency (left) and burst size (right). (**D**) Scatterplot showing the average on time ($1/k_{off}$) versus off time ($1/k_{on}$) of each gene. (**E**) Scatterplot showing the burst frequency compared to gene length. (**F**) Scatterplot showing the burst size compared to gene length. (**G**) Average burst frequency versus gene length for TOP2 inhibition affected genes (Fold Change > 1.5 and FDR < 5 x 10^-5). (**H**) Boxplot showing the burst frequency of TOP2 inhibition affected genes (Fold Change > 1.5 and FDR < 5 x 10^-5). (**I**) the same as (**H**) but with cutoff (Fold Change > 1.5 and FDR < 0.01).

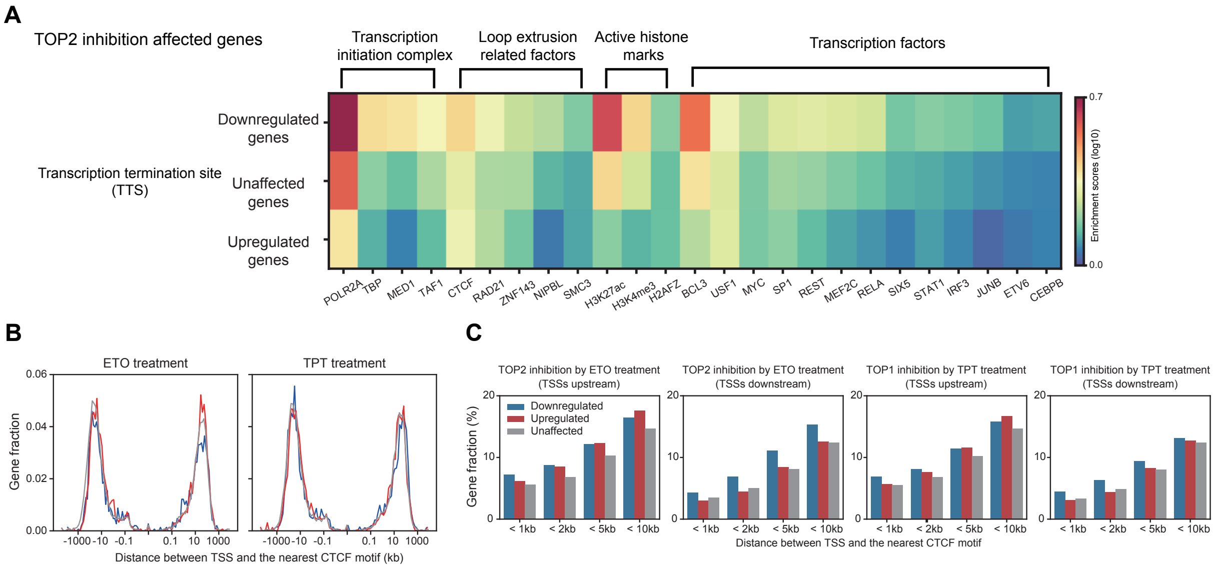


**Fig. S16. Features of genes related to loop extrusion process.** (**A**) Protein enrichment features of each group of genes affected by TOP2-inhibition. (**B**) Frequency distribution of the nearest CTCF to gene promoter, genes affected by TOP2 inhibition (left) and TOP1 inhibition (right). (**C**) Histogram of genes with CTCF binding within 1 kb, 2 kb, 5 kb and 10 kb, upstream and downstream are separated.
